# WNT inhibition primes the transcriptional landscape of mesoderm to initiate a phased ventricular cardiomyocyte specification programme

**DOI:** 10.1101/2025.11.11.687613

**Authors:** V Velecela, A Bassil, E Fawcett, E Smith, D Konstantopoulos, F Salmén, AS Bernardo, S Hoppler

## Abstract

**Background and Aims:** Cardiac reprogramming holds promise for treating ischemic heart diseases and advancing personalised medicine, but current approaches do not yet reliably generate human functional, homogeneous cardiomyocyte cultures. This limited success likely reflects our incomplete understanding of hierarchical transcriptional programmes guiding cardiomyocyte specification. Here, we modelled cardiomyocyte differentiation aiming to uncover gene regulatory networks (GRNs) guiding early human ventricular cardiomyocyte development. Our study focused on defining how inhibition of WNT signalling remodels the transcriptional landscape underlying human cardiomyocyte differentiation, since both *in vivo* cardiac development and *in vitro* cardiomyocyte differentiation protocols require WNT inhibition.

**Methods:** We modelled left ventricular cardiomyocyte differentiation from human pluripotent stem cells and experimentally manipulated WNT signalling to uncover transcriptional responses using single-cell RNA sequencing. Bioinformatics analysis defined cell identities, reconstructed differentiation trajectories, and inferred WNT inhibition-dependent gene expression and regulatory networks driving cardiomyocyte specification.

**Results:** We found that WNT inhibition (WNTi) decisively redirects mesoderm cells towards a cardiomyocyte progenitor fate, expanding their numbers while limiting alternative trajectories. GRN inference revealed both WNTi-dependent and -independent programmes and a hierarchical cascade of transcription factors driving the mesoderm-to-cardiomyocyte progenitor transition. Notably, *MEIS2* emerged as a central WNTi-independent regulator, while WNTi-responsive networks featured early (*ISL1*, *PBX3*, *TBX5* and *KLF1*) and late (MEF2-related genes, GATA-related genes, *PBX1*, *CREM*, *FOXP1* and *NKX3*-1) factors. Mesodermal GRNs were primed for cardiomyocyte specification in WNTi-treated cultures but, in the absence of WNTi, mesodermal GRNs remained ambiguous, activating *ISL1* and *PBX3* but failing to establish cardiomyocyte commitment and subsequent differentiation into contractile cells.

**Conclusion:** This work provides the first comprehensive dissection of WNTi-dependent and -independent regulatory hierarchies guiding human ventricular cardiomyocyte specification and highlights new transcriptional players which could improve cardiac reprogramming efficiency and fidelity.

**Translational Perspective:** Deciphering the transcriptional programmes that drive early cardiomyocyte specification has clear translational potential for regenerative therapies and cardiac reprogramming. By modelling left ventricular cardiomyocyte differentiation from hPSCs, our study highlights how inhibition of WNT signalling promotes ventricular cardiomyocyte progenitor commitment while restricting alternative fates. The discovery of both WNTi-dependent and -independent transcriptional activators, including some not previously linked to cardiomyocyte development, provides new insights for improving the efficiency and fidelity of direct cardiac reprogramming. Importantly, these TF candidates may also guide the development of targeted therapies for ischemic heart disease and inform personalised approaches for repairing and regenerating damaged myocardium.

## INTRODUCTION

The heart is the first organ to develop but rapidly loses its regenerative capacity after birth^1^. The mortality rate for cardiovascular disease (CVD) remains high, especially for myocardial ischemic injuries like myocardial infarction or heart failure^2^. Regenerative medicine approaches are being investigated including cell therapy and cardiac reprogramming^3^. Cardiac reprogramming is an exciting approach, which has succeeded to treat the ischemic myocardium in animal models^4^.

Cardiac reprogramming was first accomplished in cultured mouse fibroblasts using three transcription factors (TFs): *Gata4*, Mef2c and *Tbx5* (GMT)^5, 6^. Despite the high reproducibility of this approach in mice, it is not directly applicable to human cells^6^, underscoring key interspecies differences. Approaches to reprogram human cells into cardiomyocytes are different in important ways. Some use GMT plus *MYOCD* and/or micro-RNAs, small molecules or protein ligands, while others do not use *MEF2C* at all^6^. A completely new cocktail without GMT has been proposed consisting of *MYOCD*, *SMAD6* and *TBX20*^7^. However, none of these approaches reliably generates functional and homogeneous cardiomyocyte cultures: many cardiomyocyte-like cells fail to contract, cultures consist of mixed ventricle, atrial and nodal subtypes, and reprogramming efficiency remains low (5-40%)^6–8^. The limited success of these approaches in human, despite relying on known cardiac factors, likely reflects our incomplete understanding of hierarchical transcriptional programmes, constraining efforts to faithfully reproduce cardiomyocyte reprogramming in vitro. Advancing the field will require deeper insight into human cardiomyocyte specification.

Human pluripotent stem cells (hPSCs) are a powerful model for studying human cardiovascular development^9^, and successful protocols have been established for differentiating hPSCs into cardiomyocytes, generating mixed or enriched atrial-or ventricular-cardiomyocyte populations^10–14^. All such protocols rely on WNT signalling modulation, a key signalling pathway which regulates embryonic heart development^15, 16^.

The functional role of WNT signalling during heart development is complex since it promotes and restricts myocardial differentiation and is involved in cardiomyocyte expansion. Specifically, WNT pathway activation is first required for mesoderm specification^17^, while cardiomyocyte progenitor differentiation relies on inhibition of WNT signalling^18, 19^, and cardiomyocyte expansion relies again on WNT activation^20^. Mainly canonical WNT signalling has been explored in the context of heart development, however both canonical and non-canonical WNT signalling are dynamically activated throughout cardiomyocyte differentiation^21^. Moreover, WNT signalling inhibition (WNTi) was proposed to act as a ‘roadblock remover’ by downregulating the expression of CDX1/2 and MSX1 to enable cardiomyocyte progenitor specification^22, 23^. However, WNTi must play a more complex role, since even when these proposed roadblock genes are knocked-down, cardiomyocyte differentiation still requires experimental WNTi. Investigating the transition from mesoderm to cardiomyocyte progenitors in a human cell model will further our understanding of heart development and may hold the key to improving cardiac reprogramming.

Here we investigated how WNT signalling inhibition guides ventricular cardiomyocyte progenitor specification by using hPSCs and following a left ventricle-specific differentiation protocol^13^. Through single-cell transcriptomics and bioinformatic inference of Gene Regulatory Networks (GRNs), we identified how WNT signalling inhibition reshapes the transcriptional landscape of the mesoderm, progressively activating a cardiomyocyte transcriptional programme with distinct early and late phases. Our analysis reveals key transcriptional regulators involved in this mesoderm-to-cardiomyocyte progenitor transition, including cardiomyogenic factors that while active remain insufficient to drive cardiomyocyte commitment in the absence of WNTi.

## METHODS

### Human pluripotent stem cell lines: Origin, characterization and maintenance

All hESC/hiPSC experiments were approved by the UK Stem Cell Bank Steering Committee. Feeder-free H7 hESCs (WA07; 46,XX; WiCell) and MYL2-GFP hiPSCs (AICS0060-027; 46,XY; Allen Institute) were used. H7 was maintained on growth factor–reduced Matrigel (Corning); MYL2-GFP on Geltrex (Thermo Fisher). Both were cultured in mTeSR1 medium (STEMCELL Technologies). Cultures were passaged every 3–4 days at 1:6 as aggregates using Versene (Thermo Fisher Scientific). (see also Supplementary)

### Differentiation of human pluripotent stem cell lines to cardiomyocytes

Once confluent, cells were dissociated with TrypLE Express, resuspended in mTeSR1 containing Y-27632 (Tocris), counted (NC200, Chemometec), and seeded at 0.65×10^5^ cells/cm² (H7) or 0.45×10^5^ cells/cm² (MYL2-GFP). Cells were grown for 3 days, then cardiomyocyte differentiation was performed as described previously^13^ with some modifications. (see also Supplementary)

### Immunocytochemistry

Cells were seeded on Geltrex-coated 12-well chamber slides (Ibidi) or 96-well black chimney plates (Greiner) and differentiated as described. At defined time points, cells were fixed and immunostained as previously^13^ using the antibodies listed in the Supplementary immunostaining tables. (see also Supplementary)

### Flow cytometry

Differentiated cells were dissociated with TrypLE Express (Thermo Fisher Scientific), resuspended in RPMI/B27 ± insulin, pelleted, and fixed in 2% methanol-free formaldehyde in PBS for 20 min. Cells were permeabilised/blocked in 0.5% saponin/PBS for 15 min, incubated with primary antibodies (see Supplementary Flow Cytometry Antibody table) for 45 min at room temperature, washed three times in PBS, and either analysed immediately or stained with fluorochrome-conjugated secondary antibodies diluted in 5% donkey serum in BSA/PBS. After three PBS washes, cells were resuspended in PBS/BSA and acquired on an LSR Fortessa or Symphony A3 (BD Biosciences); data were analysed in FlowJo (FlowJo LLC).

### RNA extraction, cDNA synthesis and RT-pPCR

Samples were collected in RLT Plus buffer (Qiagen), snap-frozen in liquid nitrogen, and stored at −80 °C. RNA was extracted with the RNeasy Plus kit using gDNA Eliminator columns (Qiagen), cDNA was synthesised with the Maxima First Strand cDNA Synthesis Kit (Thermo Fisher Scientific), and real-time qPCR was performed with GoTaq Master Mix (Promega) using primers listed in the Supplemental Primer Table, following manufacturers’ instructions. (see also Supplementary)

### Single Cell RNA sequencing

Differentiating hPSCs (WA07, WiCell) at day 2 and day 4 were processed for single-cell RNA-seq. Cells were dissociated with TrypLE, filtered through a 35 µm strainer (Thermo Fisher), resuspended in RPMI + B27 (no insulin) with L-ascorbic acid and 10% DMSO, and frozen at −80 °C in a CoolCell LX container. After thawing, cells were resuspended in fresh RPMI + B27 (no insulin) with L-ascorbic acid, kept on ice, stained with DAPI, and live singlets (DAPI^−^) were sorted on a CytoFLEX SRT into RPMI 1640 + B27 (no insulin). Cell counts and viability were assessed by trypan blue on a Countess II. Approximately 6,000 cells per sample were loaded onto a 10x Genomics Chromium Controller; libraries were prepared with the Chromium Single Cell 3′ v3 kit, pooled, and sequenced on a NextSeq 500. (see also Supplementary)

### Sequencing and processing with Cellranger

Raw reads were processed with Cell Ranger^24^. BCL files were converted to FASTQ files with mkfastq, then cellranger count aligned reads to GRCh28 and generated filtered feature–barcode matrices (gene-by-cell counts). Each sample yielded 2,656– 4,099 cells with ∼3.7k genes per cell.

### Sample integration and clustering

Samples imported and processed using Seurat^25^. Dimensionality reduction (PCA, UMAP) and clustering were performed; multiple resolutions were tested, and 0.5 was selected as best matching expected cell types. (see also Supplementary)

### Analysis of marker gene expression

Cluster marker genes were identified with Seurat’s FindAllMarkers. Pairwise FindMarkers tests within cell types and across samples detected stage-and treatment-dependent DE genes. Genes with FDR < 0.05 underwent GO enrichment (biological process, molecular function, cellular component) using clusterProfiler (R). UpSet plots were generated with UpSetR^26^.

### Cell-lineage trajectory analysis with Monocle pseudotime and RNA velocity

Pseudotime analysis was performed by converting the Seurat object to Monocle3^27^, which uses graph-based ordering of cells based on gene expression. After rooting the trajectory in cluster 0, we inferred a pseudotime lineage reflecting the likely cellular progression.

RNA velocity was inferred by generating LOOM files with Velocyto and analysing them in scVelo. Velocity estimates used spliced-to-unspliced transcript ratios per gene per cell to infer transcriptional activity and its direction. Seurat UMAP embeddings were imported into scVelo for consistent visualisation.

### GSEA analysis

Genes were ranked by log2-fold change (Seurat FindMarkers) against all other cells, and GSEA was run with clusterProfiler’s gseGO^28^ (Biological Process) to identify positively and negatively enriched GO terms. Results were visualised with enrichplot package.

### Cell identity analysis – Capybara

Mouse data^29^ were loaded with the “MouseGastrulationData” R software package (https://github.com/MarioniLab/MouseGastrulationData) and filtered to include E7.5, E7.75 and E8 stages. Mouse genes were mapped to human homologues using “biomaRt” library^30^, allowing only one-to-one mappings, and the humanized counts were processed to create a Seurat object, which was normalized, and scaled. Capybara^31^ was run with default settings following the tutorial from step 2 (https://www.youtube.com/@morrislabtutorials5036/featured), (see also Supplementary).

### Regulon analysis with SCENIC

We applied SCENIC (Single-Cell Regulatory Network Inference and Clustering^32^) with pySCENIC^33^ to: (1) the integrated dataset across all conditions, (2) each condition separately (D2, D4_0i_, D4_12i_, and D4_48i_), and (3) the combined WNTi-treated samples (D4_12i_ + D4_48i_). We performed two analyses: (i) identification of positive-only TF regulons (“onlypos”; TF+); and (ii) identification of TF(+) and TF(–) regulons, representing activation and repression, respectively (see Supplementary). The latter allowed the identification of dual-sign regulons.

### Inference of GRNs

SCENIC-defined TF regulons were used to infer higher-order interactions and filtered by cluster specificity; Regulon Specificity Score (RSS). For each regulon, we computed the fractions of target genes up-or down-regulated (pos/neg scores) upon WNTi and combined them into a Z-score ranging from −1 to 1, termed WNT inhibition responsive score (WRS). Specifically, first WNTi responses for each target were quantified by differential expression (Seurat’s default Wilcoxon test) between WNTi-treated (D4_12i_, D4_48i_) and control (D40i) samples in clusters 0, 1, 2, 3, 5, and 7; gene scoring increased with WNTi as positive and decreased as negative. Next, for each regulon we computed the WRS based on this combined response to WNTi of the relevant target gene set. GRNs were visualised with igraph^34^, Matplotlib^35^ or NetworkX^36^, and network comparisons were plotted with Seaborn^37^. (see also Supplementary)

## RESULTS

### WNT Signalling Inhibition Directs Differentiation Towards a Cardiomyocyte Progenitor Fate and Away from Cardiovascular Alternatives

Without WNTi cardiomyocyte differentiation is impaired^18, 19, 22^. Using a protocol which generates mesoderm cells destined to differentiate into left ventricular cardiomyocytes upon WNTi^13^, we modulated experimental WNTi between days 2-4 of differentiation by either treating cells with a WNTi continuously for 2 days (D4_48i_, long WNTi) or for the final 12 hours (D4_12i_, short WNTi), or not at all (D4_0i_; control) (Figure 1A). Both short and long WNTi induced cardiomyocyte differentiation as evidenced by flow cytometry and presence of beating cells (Figure S1A), consistent with previous reports^22^.

**FIGURE 1:**
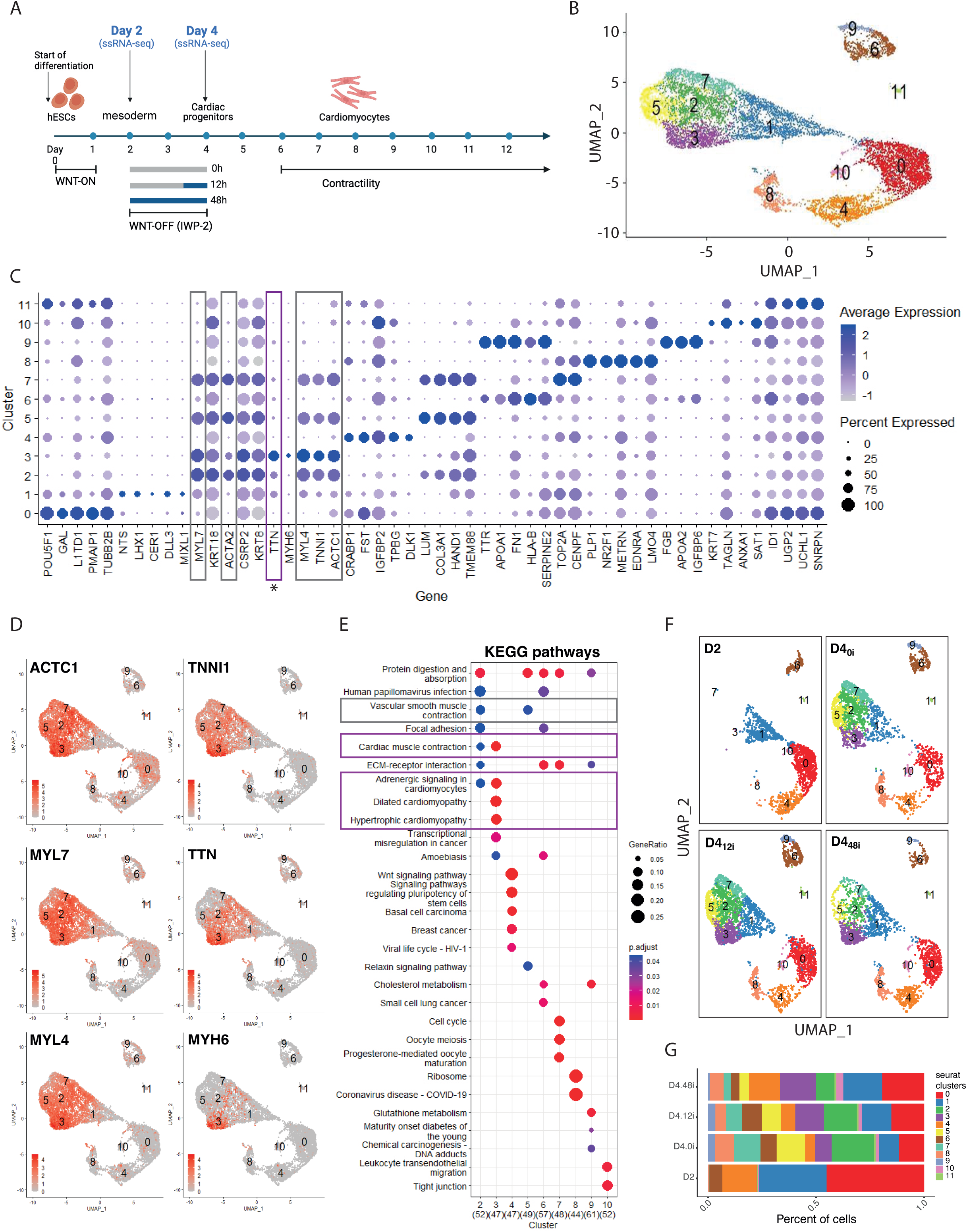
Inhibition of WNT signalling directs differentiation towards cardiomyocyte progenitors and away from alternative cardiovascular fates. **A** Experimental Design: WNT-ON = treatment with WNT agonist (CHIR99021); WNT-OFF = treatment with a WNTi (IWP-2) for 12h or 48h. Single-cell RNA-seq collection points indicated. **B** UMAP of all samples showing 12 clusters (0-11) identified. **C** Dot plot of top five markers per cluster. Clusters 2, 3, 5, and 7 share genes (grey boxes); cluster 3 is distinct with lower *ACTA2* and higher *TNN*/*MYH6* expression (purple box). D UMAP of cardiovascular marker expression (all samples combined). E KEGG pathways summary: cluster 3 enriched for cardiomyocyte terms (purple boxes); clusters 2 and 5 for vascular smooth muscle contraction (grey box). F UMAP by day **(D)** and WNTi treatment. Day 4 samples were treated with short (D4_12i_), long (D4_48i_) or no (D4_0i_) WNTi. G Cell proportion per sample per cluster.

Single-cell RNA-sequencing (days 2 and 4) identified 12 clusters (0-11, Figure 1B). Based on marker gene enrichment per cluster we inferred putative cell identities (Figure 1C and Supplemental Table 1). Clusters 2, 3, 5 and 7 were identified as cardiovascular populations, co-expressing myosin, troponin and actin filament genes (Figure 1C and Supplemental Table 1). Uniform Manifold Approximation and Projections (UMAPs) of lineage-specific markers (*ACTC1*, *MYL7*, *MYL4*, *MYH6* and *TNNI1*) supported this assignment (Figure 1D). Interestingly, cluster 3 was distinct, with lower ACTA2 (smooth muscle marker^38, 39^) and enrichment in the cardiomyocyte marker *TTN* (Figure 1C-D). Among the cardiovascular populations, only clusters 3 and 5 were in the G_1_ phase of the cell cycle, consistent with differentiation^40^ (Figure S1B).

Other identities were also identified: cluster 0 (pluripotency: *POU5F1*, *SOX2*), cluster 1 (mesoderm: *MESP1*, *EOMES*), clusters 6 and 9 (endoderm: *FOXA1*, *SOX17*), and clusters 4 and 8 (neural and paraxial mesoderm: *PAX3*, *NR2F1*, *SOX2*) (Figure S1C). Clusters 10 and 11 were small, poorly defined and excluded (Supplemental Table 1).

Upset plot analysis confirmed similarity among cardiovascular clusters and the distinctiveness of cluster 3 (Figure S1D). KEGG enrichment further identified cluster 3 as cardiomyocyte progenitors (enriched in cardiomyocyte-related terms), and distinguished it from the other cardiovascular clusters, with clusters 2 and 5 enriched for vascular smooth muscle contraction (Figure 1E). To probe deeper into the cell identities, we used Capybara^31^ to map our cells to the transcriptional labels defined in a gastrulating mouse dataset^29^, using human orthologues for cross-species comparison. Cluster 3 was readily identified as cardiomyocytes, with other cardiovascular clusters identified as mesenchyme and cluster 1 as mesoderm (Figure S1E).

Given WNTi’s role in promoting cardiomyocyte differentiation, we tested its impact on cluster proportions (Figure 1F-G). Day-4 transcriptional landscapes were remarkably similar, irrespective of WNTi treatment (Figure 1F), but WNTi increased cardiomyocyte progenitors (cluster 3) and reduced other cardiovascular clusters (2, 5, 7) (Figure 1G). Capybara confirmed cardiomyocyte enrichment with WNTi (Figure S1F). Longer WNTi (D4_48i_) further raised *NKX2*-5 and *ISL1*-expressing cardiac progenitors, while reducing *LUM-*and *COL3A1*-expressing non-cardiomyocyte cardiovascular cells (Figures S1G-H).

These differences correlate cell lineage outcomes with the duration of WNTi treatment, validating the role of WNTi in directing differentiation towards the cardiomyocyte lineage, and away from alternative cardiovascular cell fates.

### WNTi Promotes Direct Cardiomyocyte Progenitor Differentiation and Structural Gene Expression

We used RNA velocity^41^ to infer how WNTi directs cell lineage differentiation trajectories. Unrooted lineage streams (Figure 2A) confirmed the pluripotency cluster (cluster 0) as the earliest transcriptional state (lineage origin) followed by an initial transition to mesoderm (cluster 1), and subsequent divergence into cardiovascular cell fates (clusters 2, 3, 5, 7). Importantly, without experimental WNTi (D4_0i_), fewer lineage streams converged towards the cardiomyocyte progenitor cluster (cluster 3), whereas increasing WNTi (D4_12i_ and D4_48i_) revealed a trajectory suggesting direct differentiation from mesoderm (cluster 1) to cardiomyocyte progenitors (cluster 3).

**FIGURE 2:**
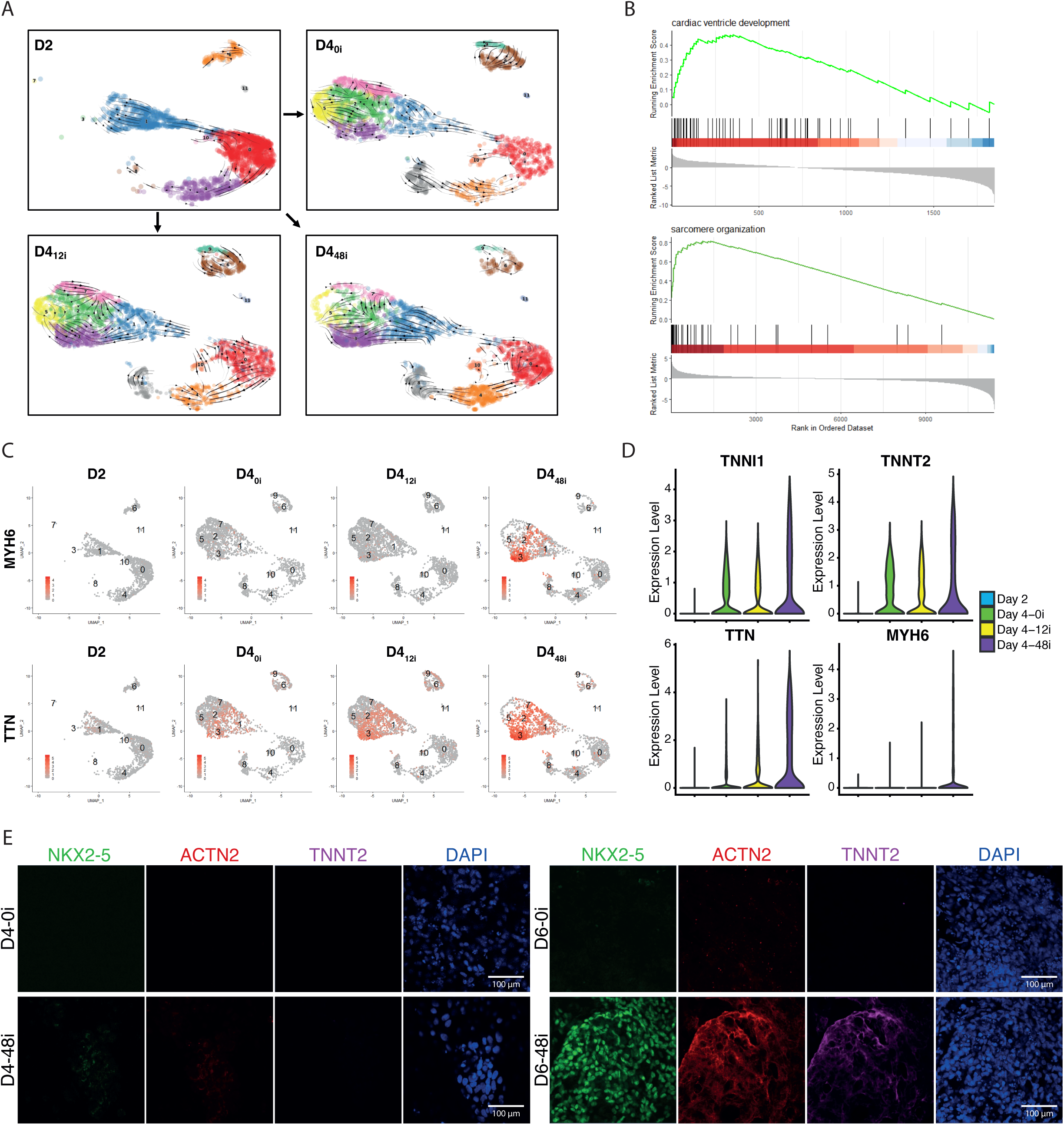
WNTi Promotes Direct Cardiomyocyte Progenitor Differentiation and Structural Gene Expression. **A** RNA velocity inferred cell lineage. **B** GSEA NES plots showing terms enriched in cluster 3. **C** UMAP of cardiomyocyte markers in day 2 samples or day 4 samples treated with short (D4_12i_), long (D4_48i_) or no (D4_0i_) WNTi. **D** Violin plots of cardiomyocyte marker expression in cluster 3. **E** Immunofluorescence of cardiomyocyte structural (TNNT2, ACTN2) and TF (NKX2-5) proteins in day (D) 4 (left) and day 6 (right) cultures with and without WNTi (48i and 0i, respectively). Nuclei were counterstained with DAPI.

Even without experimental WNTi (D4_0i_ control) some cells still acquire cardiomyocyte progenitor identity (Figures 1F-G and 2A). We therefore considered whether endogenous inhibition of WNT signalling occurs in control cells (D4_0i_). Indeed, we detected expression of known endogenous WNT inhibitors *SFRP5*^42^ and *TMEM88*^43^ within cluster 3 (Figure S2A) and independently confirmed their increasing expression after mesoderm induction (Figure S2B).

To better characterise cardiomyocyte progenitor cluster 3, GO term analysis was used, revealing enrichment for cardiac muscle development, differentiation and morphogenesis, as well as contraction and sarcomere terms (Figure S2C). Gene set enrichment analysis (GSEA) corroborated this and further highlighted an enrichment for ventricular cardiac muscle tissue terms, consistent with our ventricular protocol^13^, while negative normalised enrichment scores (NES) confirmed the reduced proliferative capacity of this cluster (Supplemental Table 2, Figure 2B and S2D). Key cardiomyocyte structural genes (*TTN*, *MYL4*, *MYH6*, *TNNI1* and *TNNT2*) were associated with this cluster with their expression increasing with prolonged WNTi (Figures 2C-D and Supplemental Table 1). Notably, as proteins these markers were only detected at the onset of beating; e.g., TNNT2 and ACTN2 proteins were first detected at day 6 of differentiation in WNTi-treated cultures (Figure 2E), i.e. two days post RNA detection.

Together our results indicate that WNTi promotes not only mesoderm specification towards cardiomyocyte progenitors but also primes these progenitors to express structural contractility proteins.

### WNTi Modifies the Transcriptional Landscape of Cardiomyocyte Progenitor Cells

To probe how WNTi promotes cardiomyocyte differentiation, we assessed TFs up-or down-regulated in cluster 3 following WNTi. We defined “early” responders as TFs altered after short WNTi (D4_12i_) and “late” responders, potentially forming part of a secondary response, as TFs unique to long WNTi (D4_48i_); TFs shared between D4_12i_ and D4_48i_ were considered reinforcers of a sustained WNTi response. We first performed differential expression analysis between controls (D4_0i_) and both WNTi-treated samples (D4_12i_ and D4_48i_) (Figure 3A and Supplemental Table 3). We then temporally mapped these WNTi-responsive TFs along a 60-day differentiation timeline using a published bulk RNA-seq dataset^13^, validating genes that peaked before (e.g. MXS1, CDX1, CDX2) or during the WNTi step (e.g. PBX3, ISL1 and ZNF608) as candidates for cardiomyocyte specification (Figure S3A).

**FIGURE 3:**
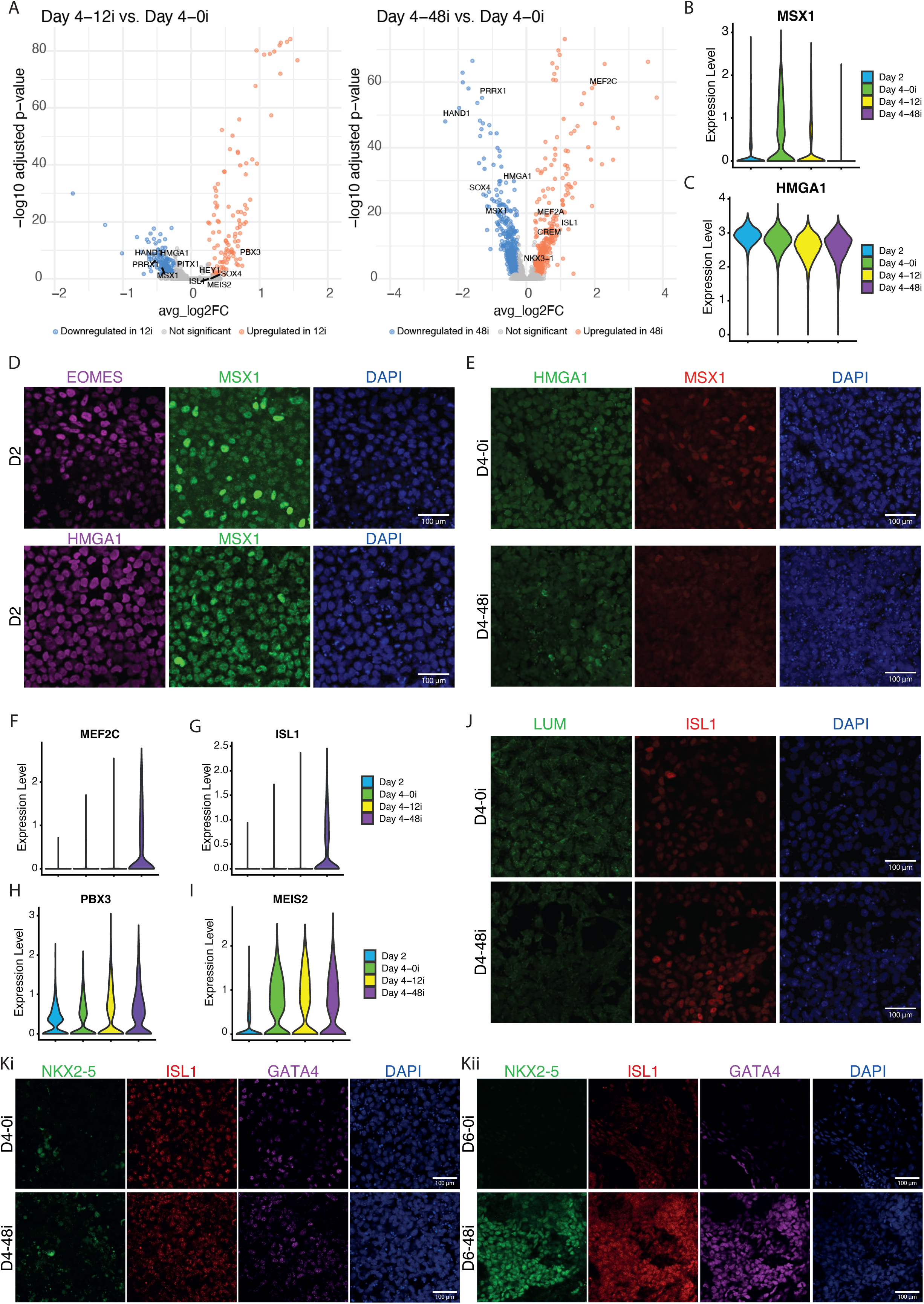
WNTi Modifies the Transcriptional Landscape of Cardiac Progenitor Cells. **A** Volcano plot of differential expression comparing WNTi-treated (D4_12i_, D4_48i_) vs control (no-WNTi, D4_0i_). Top TFs highlighted. **B-C** Violin plots of *MSX1* (**B**) and HMGA1 (**C**) expression. **D-E** Immunofluorescence for MSX1, HMGA1 and EOMES at day 2 (**D**) and day 4 (**E**) cultures grown with (D4_48i_) or without WNTi (D4_0i_). Nuclei were counterstained with DAPI. **F-I** Violin plots of cardiomyocyte TF expression in cluster 3. **J-K** Immunofluorescence for LUM and ISL1 (**J**) or NKX2-5, ISL1 and GATA4 expression (**K**) at days 4 or 6 with (D4_48i_, or D6_48i_) or without WNTi (D4_0i_, D6_0i_). Nuclei were counterstained with DAPI.

It has previously been shown that WNTi removes ‘roadblocks’ to cardiomyocyte progenitor specification^22^. We therefore examined TFs downregulated by WNTi in cluster 3. Of the reported ‘cardiac roadblock’ TFs^22^, only MSX1, but not CDX1 or CDX2, responded to WNTi in our results (Supplemental Table 3, and Figures 3B and S3B-D). Notably, MSX1^+^ cells were rare in cluster 3 even without WNTi, whereas MSX1 was abundant in other cardiovascular clusters (2, 5 and 7) (Figure S3B), consistent with a role in blocking cardiomyocyte progenitor differentiation within those clusters.

Interestingly, several genes not previously linked with WNTi were downregulated as early responders (Supplemental Table 3, and Figure 3A). However, only High Mobility Group AT-Hook 1 (*HMGA1*) was highly expressed at day 2 (Figure S3A) and significantly reduced after short WNTi (D4_12i_) (Supplemental Table 3, and Figure 3B). HMGA1 is a structural protein that regulates transcription by changing the conformation of DNA, which can amplify WNT signalling^44^. We confirmed the presence of MSX1 and HMGA1 proteins at day 2 (Figure 3D) and within D4_0i_ samples. MXS1 decreased with WNTi at day 4, whereas HMGA1 expression was similar between D4_0i_ and D4_12i_ but showed reduced nuclear and increased cytoplasmic staining after D4_48i_ (Figure 3E and S3C). These patterns suggest that lowering expression of these genes (and/or nuclear loss for HMGA1) is required for cardiomyocyte differentiation.

We next examined TFs upregulated following WNTi in cluster 3 (Figure 3A and Supplementary Table 3). These included: *PBX3*, a gene associated with congenital heart disease^45^; *MEF2C*, a key regulator of cardiovascular development widely used in cardiac reprogramming^5^; and *ISL1*, a marker of undifferentiated cardiomyocyte progenitors that is a key player in cardiac development^46^ (Figure 3F-H and S1H). Among the short WNTi-only upregulated genes was *MEIS2* (Figure 3I), known to drive hPSCs differentiation towards cardiovascular cell types^47^ and to interact with PBX genes to regulate transcription^48^, though it was also expressed in other cardiovascular clusters (Figure S5Gi). We confirmed presence of ISL1 protein at day 4, but its expression was similar in control (D4_0i_) and WNTi-treated samples, despite a trend for more ISL1^+^ cells with longer WNTi and elevated *ISL1* transcript in D4_48i_ (Figure 3G, 3J-K and S3D-E). ISL1 co-localised with the non-cardiomycyte cardiovascular progenitor marker LUM at day 4 (Figure 3J) and with the cardiac progenitor marker NKX2-5^49^ at day 6 (Figure 3K), suggesting ISL1 is a broad cardiovascular marker early in development. Likewise, GATA4, another cardiac development-associated TF^50, 51^, was co-expressed with some ISL1^+^ cells, regardless of WNTi treatment, but both ISL1 and GATA4 increased by day 6 with WNTi, while little to none remained without WNTi (Figure 3K).

Overall, these results suggest that, beyond removing any roadblock, WNTi promotes activation of a cardiomyocyte-specific transcriptional programme, which preserves ISL1 and GATA4 expression and activates NKX2-5, enabling subsequent structural protein expression.

### WNTi Restricts Expression of Repressor **TFs** Revealing Novel Inhibitors of Cardiomyocyte Differentiation

We used SCENIC^32, 33^ to refine our transcriptomic analysis and identify TFs acting as repressors of cardiomyocyte differentiation. This revealed negative regulons: i.e. transcriptional repressors whose reduced expression in our samples facilitates de-repression and therefore upregulation of relevant associated target genes. Regulon activity was scored based on the number of genes potentially regulated within each cluster and we calculated a regulon specificity score (RSS) per cluster (Supplemental Table 4).

Day-4 negative regulons differed between controls (D4_0i_) and WNTi (D4_12i_, D4_48i_) (Supplemental Table 4, and Figures 4A and S4A-B). Reassuringly the pluripotency cluster (0) was enriched for negative regulons headed by mesodermal TFs (*SNAI2*^52^) and cardiomyocyte lineage specification TFs (*MEIS1*^53^ and *GATA4*^50, 51^) (Figures 4A and S4A-C). The mesoderm cluster (1) shared negative regulon activity with clusters 2, 3, 5 and 7 (Figures 4A and S4A-C), but lacked specific enrichment, confirming its intermediate state.

**FIGURE 4:**
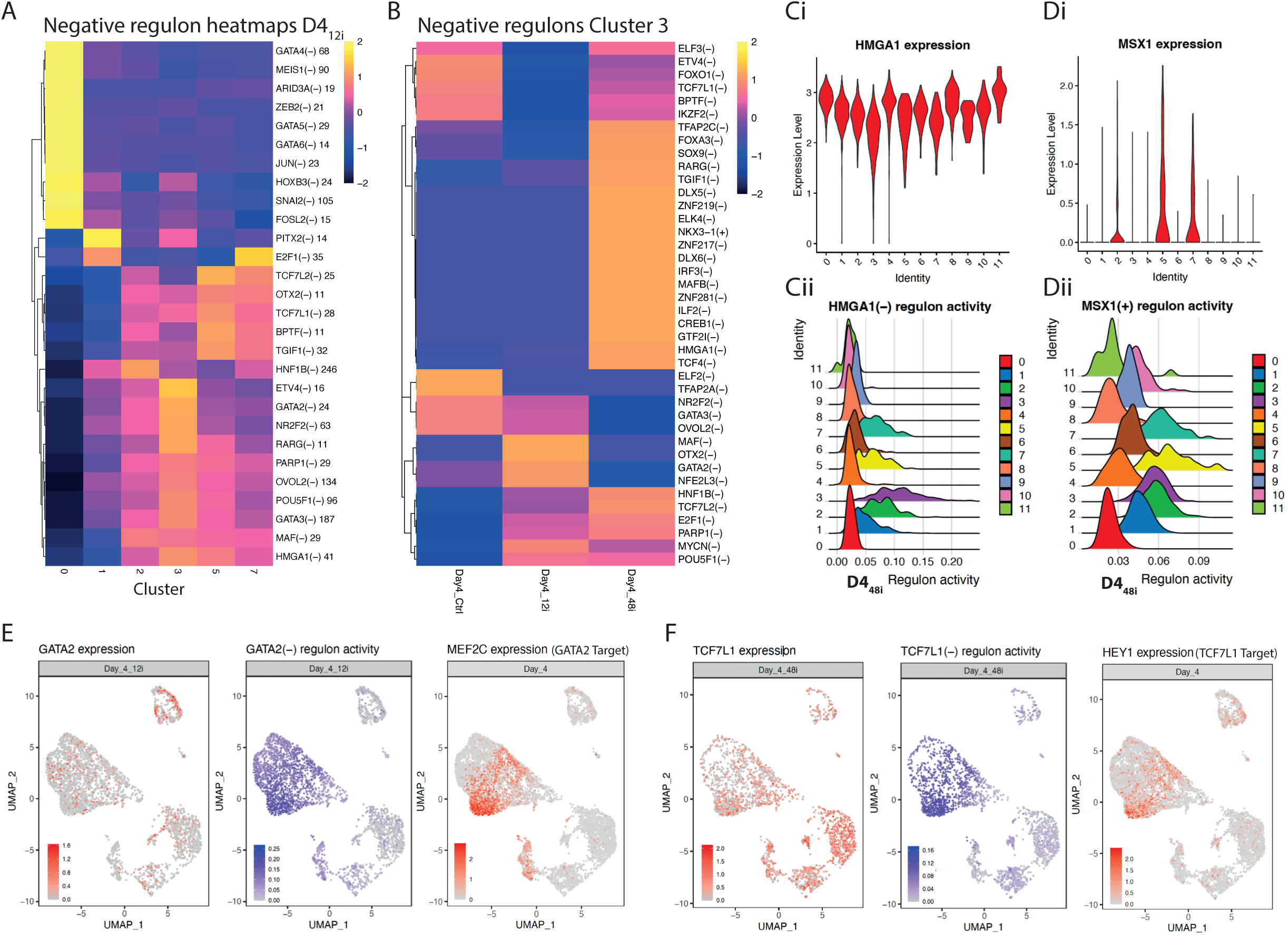
WNTi Restricts Expression of Repressor TFs, Revealing Novel Inhibitors of Cardiomyocyte Differentiation. **A** Heatmap of negative regulon activity across indicated clusters in day (D) 4 samples treated with short WNTi (D4_12i_). **B** Heatmap of negative regulon activity in cluster 3 (day 4) with short (D4_12i_), long (D4_48i_) or no (D4_0i_) WNTi. **C** *HMG1A* expression (**Ci**: violin plot) vs HMGA1(-) regulon activity (**Cii**: regulon activity ridge plot) across clusters. **D** *MSX1* expression (**Di**) vs MSX1(+) regulon activity (**Dii**) across clusters. **E** UMAPs comparing *GATA2* expression with GATA2(-) regulon activity and with *MEF2C* expression (negative GATA2 target). **F** UMAPs comparing *TCF7L1* expression with TCF7L1(-) regulon activity and *HEY1* expression (negative TCF7L1 target).

In cluster 3, among its downregulated TFs (Figure 3A and Supplemental Table 3), only *HMGA1* directed a cardiovascular-specific negative regulon, enriched after long WNTi (D4_48i_) despite widespread *HMGA1* expression (Supplemental Tables 3-4, Figures 3E, 4A-C, S3C and S4B). Contrary to expectations^22^, *MXS1* did not form a negative regulon (Supplemental Table 4), instead it broadly directed a positive regulon (i.e. potentially acting as a transcriptional activator) in cardiovascular clusters, including in cardiomyocyte progenitors (cluster 3) after long WNTi (D4_48i_), despite low expression (Figures 3E, 4D and S3C). Additional negative regulons in cardiomyocyte progenitors often strengthen by WNTi included: *POU5F1* (a key pluripotency marker^54^) activated following short or long WNTi; negative regulons with enriched activity in D4_12i_ like *NR2F2* (typically associated with atrial cardiomyocyte differentiation^55^), *OVOL2* (considered a repressor of epithelial-mesenchymal transition^56^), and *GATA2* (known to promote hematopoietic development and repress cardiac differentiation^57^); and others with enriched activity after long WNTi such as *RARG* (a retinoic acid receptor known to restrict the size of the first heart field^58^), and *TCF7L1* (a transcriptional repressor in the absence of WNT signalling^59^ known to restrict cardiomyocyte differentiation and promote endothelial specification^60^) (Supplemental Table 4, Figures 4B, 4E-F and S4D-E). De-repressed targets of *GATA2* and *TCFL71* included cardiomyocyte development-linked TFs like *MEF2C* and *HEY1*^61^, respectively (Supplemental Table 4 and Figures 4E-F). Interestingly, several negative regulons (*HMGA1*, *NR2F2*, *GATA2* and *TCF7L1*) also appeared in D4_0i_ (control) cluster 3, but with lower RSS (Supplemental Table 4 and Figure S4D).

Together this analysis shows that WNTi reduces expression of TFs (negative regulon TF heads) that repress cardiomyocyte differentiation and favour alternative lineages, thereby de-repressing their targets (negative regulon target genes), especially in cardiomyocyte progenitors.

### WNTi Activates Expression of TFs Promoting Cardiac Specification, Revealing Novel Drivers of Cardiomyocyte Differentiation

To identify TF activators downstream of WNTi, we reapplied SCENIC to call positive regulons, i.e. TFs whose increased expression correlates with an upregulation of downstream target genes (Figures 5A and S5A-B and Supplemental Table 4). Reassuringly, the pluripotency cluster (0) was dominated by *POU5F1* and *SOX2* positive regulons, consistent with their role in somatic-to-pluripotency reprogramming^62, 63^ (Figures 5A, S1C, and S5A-D).

**FIGURE 5:**
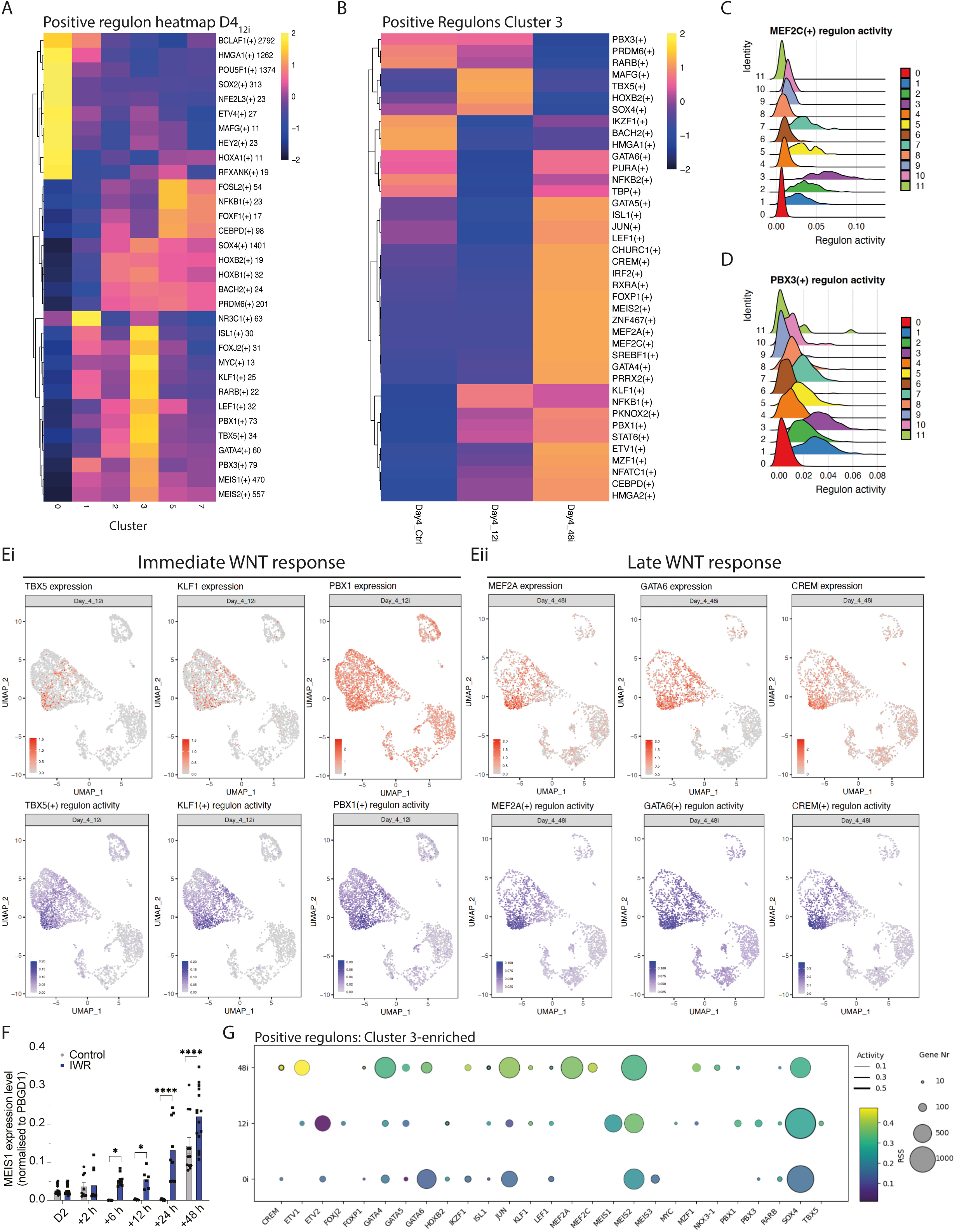
WNTi Activates Expression of TFs Promoting Cardiac Specification, Revealing Novel Drivers of Cardiomyocyte Differentiation. **A** Heatmap of positive regulon activity in day 4 samples cultured with WNTi (D4_12i_); analysed clusters indicated. **B** Heatmap of positive regulon activity in cluster 3 (Day 4) under short (Day4_12i_), long (Day4_48i_), or no (Day4_0i_) WNTi. **C-D** Regulon activity plots for MEF2C(+) (**C**) and PBX3(+) (**D**). **E** UMAPs showing TF gene expression vs associated positive regulon activity for early (D4_12i_, **Ei**) and late (D4_48i_, **Eii**) WNTi responses. **F** *MEIS1* expression time course by RT-qPCR. One-way ANOVA; mean ± s.d.; n=3. **G** Bubble plot of top positive regulons in cluster 3 (day 4) within Day4_12i_, Day4_48i_, or Day4_0i_ conditions. Regulon activity (line thickness), specificity (RSS, bubble colour), and number of predicted targets (Bubble size) are indicated.

At day 4 (even with WNTi), non-cardiomyocyte cardiovascular clusters (2, 5 and 7) were enriched for positive regulons headed by: *FOSL2* (an AP-1 TF component expressed in cardiac lineages and involved in fibrosis and vascular remodelling^64, 65^); *CEBPD* (which mediates epicardial activation during heart development and injury^66^); and the proposed cardiomyocyte ‘roadblock’ TF *MSX1*^22^ (Figures 5A, 4D, S5A-B and S5E-F). Their downstream genes (e.g. *HAND1*, *MIXL1*, *COL3A1* and *ADAM19*) were upregulated in these clusters, especially without WNTi, implying their involvement in specifying these non-cardiomyocyte cardiovascular lineages (Supplemental Table 4 and Figures S1G and S4E-F).

In cardiomyocyte progenitors (cluster 3) several WNTi-induced TFs (Figures 3 and S3) directed positive regulons, including: *PBX3*, *MEF2C*, *ISL1*, and *MEIS2* (Supplemental Table 4 and Figures 5A-D and S5G). Interestingly, *ISL1*, *MEIS2* and *PBX3*-controlled regulons were also active without WNTi. Among the early (short WNTi) top positive regulons were also regulons headed by *ISL1*, *PBX3* and *MEIS2*, along with other key cardiac regulators like: *MEIS1* (a key regulator of cardiac proliferation^53^); *TBX5* (a marker of the first heart field used in cardiac reprogramming^5^); *PBX1* (involved in distinct regulatory networks patterning the great arteries and the heart^67^); GATA4 (a well-known TF regulator of heart formation, differentiation, and hypertrophy, whose mutant phenotype is linked to various congenital heart defects in humans and mice^50, 51^); and *KLF1* (a core transcriptional regulator of cardiomyocyte renewal in adult zebrafish hearts^68^) (Figures 5B, 5D and 5Ei). Late (long WNTi) regulons were headed by: *MEF2C*^5, 61^ (Figure 5C); *MEF2A* (critical for myocardial function^69^); *GATA6* (a well-known TF regulator of cardiac-specific gene expression during embryogenesis^51^); and *CREM* (whose repressor isoforms are induced after stimulation of the β-adrenoceptor leading to arrhythmogenic alterations in ventricular cardiomyocytes^70^) (Figure 5Eii). Real time RT-PCR confirmed *MEIS1* and *PBX3* rise shortly after WNTi (Figures 5F and S5Hi). However, *ISL1* expression was upregulated regardless of WNTi (Figures 3J-K, S3D-E and S5Hii), and *MEF2C* increased only after long WNTi, confirming a late response (Figure S5Hiii). Target sets within these regulons confirmed their link to cardiomyocyte progenitor specification with multiple cardiac sarcomere genes present (Supplemental Table 4).

Deeper analysis of the most active cardiomyocyte progenitor regulons showed that activity and specificity did not necessarily correlate with the size of the regulated gene set within the regulon and was generally more specific in D4_48i_ samples and less specific in D4_0i_ samples (Figure 5G). Some regulons were WNTi-independent, including *GATA4*, *GATA5*, *ISL1*, *JUN*, *MEIS2*, *RARB* and *SOX4* regulons, whereas others were WNTi-specific, like *PBX1*, *NKX3*-1, *MEIS1*, *MEF2A* and *MEF2C*, *TBX5*, *KLF1*, *MZF1* (a gene known to modulate cardiogenesis by interacting with an *NKX2*-5 cardiac enhancer^71^), and *ETV1* (which is responsible for specification of rapid conduction zones in the heart^72^). Interestingly, several regulon heads (e.g. *ISL1*, *MEIS2*, *GATA4*, *SOX4*) were ubiquitously active regardless of WNTi despite cluster 3-specificity, suggesting these play co-factor roles rather than being primary instructors of cardiac specification (Figures S5A and S5I).

A surprising cluster-3-specific positive regulon without a WNTi correlation was headed by *LEF1* (a TCF/LEF factor that binds β-catenin to activate WNT target genes^59^) (Figure 5G). *LEF1* expression decreased with WNTi (Supplemental Table 3), consistent with it being a positive feed-back target of WNT as previously reported^73^, and suggesting its regulon activity may be a transient result of LEF1 still being expressed in the cultures, rather than playing a direct role in cardiomyocyte specification.

In all, these results indicate WNTi remodels transcription gradually: some activators are induced early and sustained, while others emerge later as a secondary response. Dissecting the gene regulatory networks (GRNs) orchestrating these steps will be essential to define the drivers of cardiomyocyte specification.

### WNTi Establishes a TF-based Gene Regulatory Network for Cardiomyocyte Specification

Combinatorial TF interactions drive context-dependent gene expression and differentiation during development^74^. Using SCENIC, we identified downstream gene sets (regulons) across all clusters and samples (Supplemental Table 4). To predict the global GRNs promoting commitment to cardiomyocyte progenitors, we inferred TF networks using GRNboost (pySCENIC)^33, 75^; analysis of the full dataset was performed to rank source-target TF interactions by importance score (IS) (Supplemental Table 5). We prioritised TFs that: (i) are expressed in cluster 3; (ii) control active regulons in cluster 3 (based on regulon specificity score, RSS); and (iii) have high likelihood source-target interactions; and (iv) regulate many TF targets (dominance within the GRN).

Reassuringly, the inferred global GRNs differed between control and WNTi conditions, and between short and long WNTi, with partial overlap of core TFs (Supplemental Table 5 and Figures 6A-C). Control (D4_0i_) networks were distinct, with *MEIS2*, *ISL1*, *PBX3* and *SOX4* being the main core TFs (TFs at the centre of the GRN) shared with WNTi core GRNs (Figures 6A-C).

**FIGURE 6:**
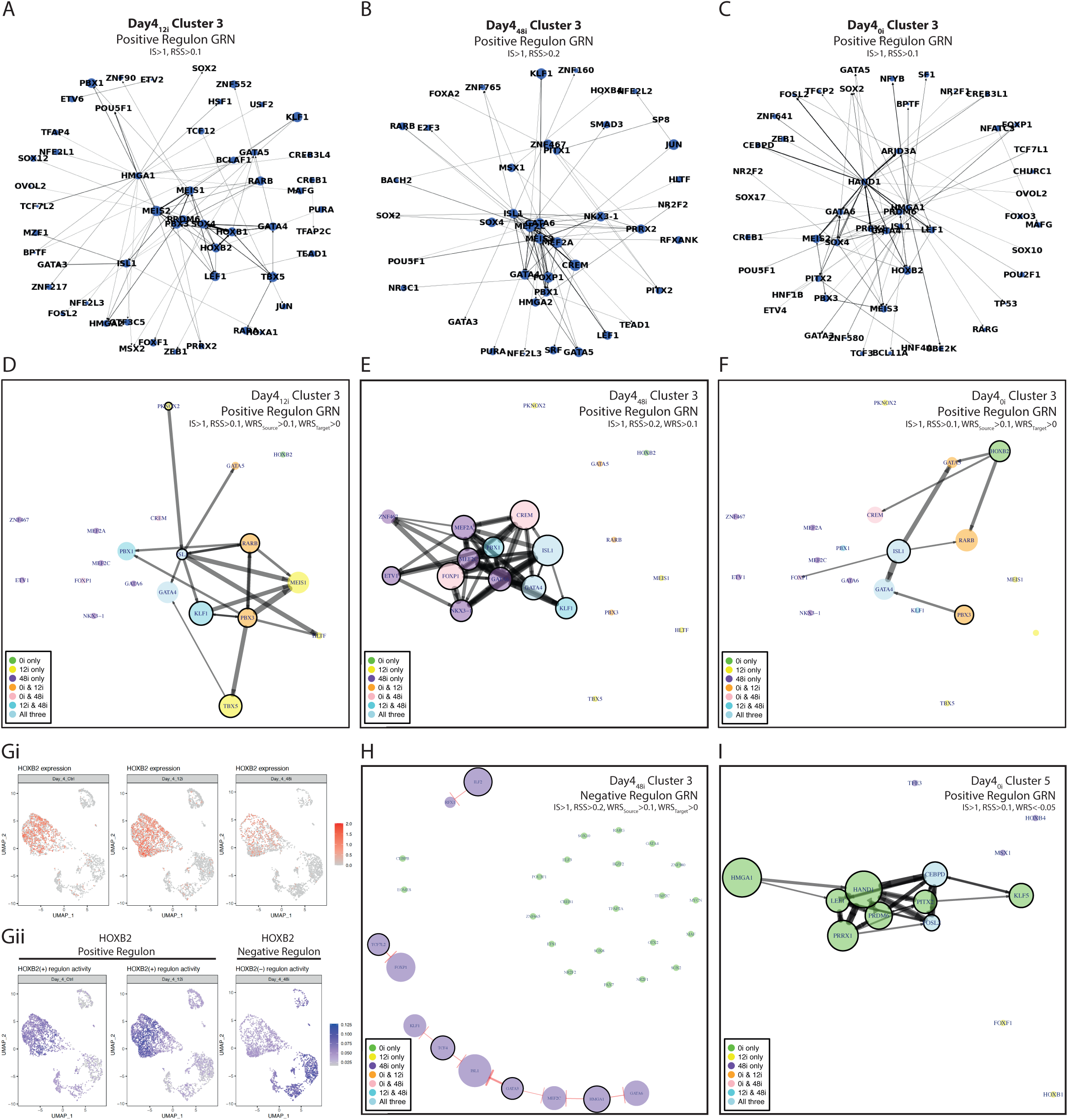
WNTi Establishes a TF Gene Regulatory Network for promoting Cardiomyocyte specification in cluster 3 A-C. Global cluster 3 TF activators GRNs (day 4) under short (Day4_12i_, **A**), long (Day4_48i_, **B**), or no (Day4_0i_, **C**) WNTi. Top 100 interactions shown (importance score, IS≥1; regulon specificity score, RSS≥0.1 for D4_0i_ and D4_12i_ or ≥0.2 for D4_48i_). **D-F** WNTi-responsive cluster 3 TF GRNs with WNTi-enabled positive regulon activity (node size) under Day4_12i_ (**D**), Day4_48i_ (**E**) or Day4_0i_ (**F**). Colour coding indicates whether a TF is active across all or only a subset of experimental samples (see legend). A border around the TF bubble denotes a source TF, while border absence indicates it functions only as a target under the defined thresholds. Line thickness reflects the importance score of each source–target interaction. **G** *HOXB2* expression (**Gi**) and regulon activity (**Gii**) across day 4 samples (Day4_12i_, Day4_48i_, and Day4_0i_). **H** Cluster 3 TF GRN of transcriptional repressors (negative regulon activity, node size) under long WNTi (Day4_48i_). **I** Cluster 5 TF GRN of WNTi-repressed positive regulon activity (node size) under no WNTi (Day4_0i_). For panels **D-I**: colours indicate TF activity across all vs subset of conditions (see legend); a node border marks a source TF (no border = target only); interaction line thickness scales with IS.

To highlight TFs and interactions particularly relevant to the WNTi response, we developed a WNTi response score (WRS) integrating how targets within each TF-controlled regulon (source) change with WNTi (see Methods and Supplemental Table 5). For the early response (D4_12i_) we used a low WRS cutoff for the targets (reflecting limited time for target activation). *ISL1*, *PBX3*, *RARB*, *KLF1* and *TBX5* emerged as main sources within the WNTi-focused GRN (Figure 6D), with *ISL1* and *PBX3* central by target number and position within the global GRN (Figure 6A). By contrast, *GATA4* and *MEIS1*, well known cardiomyocyte TFs^50, 51, 53^ and regulon heads in this condition (Figure 5G), acted primarily as targets within the WNTi-focussed analysis, indicating weak/absent response to short WNTi (Supplemental Table 5 and Figure 6D). For the late WNTi response, a stricter WRS cutoff was applied, revealing a shifted GRN. Only *ISL1*, *PBX1* and *KLF1* were shared between early and late WNTi-responsive networks, with the late GRN comprising *ISL1*, *PBX1*, GATA TFs (*GATA4* and *GATA6*) and MEF2 TFs (*MEF2A* and *MEF2C*) TFs, together with *FOXP1*, *NKX3-1* and *CREM* (Figure 6E), all of which were at the core of the global GRN (Figure 6B).

To identify factors essential for full cardiomyocyte differentiation, we compared these GRNs with the one inferred in cluster 3 without WNTi (non-beating) (Figure 6F). WNTi-treated and -untreated samples shared *GATA4* and *ISL1* (present in all Day 4 samples); *PBX3*, *RARB* and *GATA5* (shared control and short WNTi); and *CREM* and *FOXP1* (shared control and long WNTi). However, the control GRN included *HOXB2* (Figure 6F), potentially counteracting the *ISL1* initiated programme. Interestingly, *HOXB2* drove a positive regulon across cardiovascular clusters in no-/short-WNTi, but only a negative regulon after long WNTi (Supplemental Table 4 and Figures 5G and 6G), was significantly downregulated with WNTi (Figure 6G), and had a negative WRS in WNTi samples (Supplemental Table 5), supporting a repressing role for *HOXB2* in cardiomyogenesis.

Although WNTi reduces expression of the proposed roadblock TF MSX1^22^, it was not identified as negative regulon here. Instead, alternative transcriptional repressors, including HGMA1, were identified (Supplemental Table 4, and Figures 4 and S4). GRNboost further revealed negative TF interactions centred on the source TFs *HMGA1*, *GATA3*, *TCF4*, *ILF2* and *TCF7L2*, acting as transcription repressors of core cardiac progenitor TFs including: *ISL1*, *MEF2C*, *GATA6* and *KLF1* (Supplemental Table 5, Figures 6H and S6A-B). This repression became evident only after long WNTi, consistent with the time required for repressor function to take effect (Supplemental Table 5 and Figures 6H and S6A-B).

Because no cardiomyocytes arise without WNTi and non-cardiomyocyte clusters expand (Figure 1F), we interrogated GRNs in these lineages. In cluster 5, the most differentiated (Figure S1B), the GRN was distinct from that of the cardiomyocyte progenitor cluster (3) and comprised interactions between genes involved in: (i) growth and proliferation (*HMGA1*^44^, CEBPD^76^, *FOSL2*^77^ and *PRDM6*^78^); (ii) cardiac development (*HAND1*^79^ and *PITX2*^80^); (iii) WNT signalling and mesodermal development (*LEF1*^59^ and *PRRX1*^81^); and (iv) smooth muscle development (*PRDM6*^78^) (Supplemental Table 6, Figure 6I). After WNTi, *CEBPD* and *FOSL2* remained part of the GRN (Figures S5E-F) but, with long WNTi (D4_48i_), the few remaining cells induced GRN changes including the activation of *MSX1*, a known ‘cardiac roadblock’^22^ (Figure S6C-D), suggesting clear association of *CEBPD* and *FOSL2* with this cluster and possibly a *MSX1*-mediated locking of cluster 5 identity (negative correlation with WNTi).

Together these analyses show that WNTi acts as an organiser, not merely a permissive cue, of the cardiomyocyte specification GRN, suppressing alternative programmes (via novel repressors), releasing cardiac targets, and assembling a committed network consisting of classic and novel activators set in motion in a gradual manner. This GRN framework clarifies why differentiation stalls without WNTi and highlights testable levers to steer fate.

### WNTi Primes the Transcriptional Landscape of Mesoderm for Cardiomyocyte Differentiation

RNA velocity showed that WNTi drives a directional trajectory from mesoderm into cardiomyocyte progenitors (Fig. 2A). To test whether these streams were reflected in mesoderm GRNs, we analysed cluster 1 using regulon specificity filters (but no WRS filter) (Supplemental Table 7). In day 2 cultures, mesoderm identity depended on *EOMES*, *POU5F1* and *MIXL1* (canonical mesoderm TFs^82–84^) and, consistent with a cardiac-primed state, the GRN also included cardiac genes such as *GATA4*, *GATA6*, *ISL1* and *MEIS1* (Supplemental Table 7, Figure 7A). By day 4, the transcriptional landscape and GRN architecture had shifted markedly irrespective of WNTi, with core mesoderm TFs like *POU5F1* and *MIXL1* no longer prominent (Supplemental Table 7 and Figures S7A-C). Day 4 GRNs diverged by condition: with WNTi, mesoderm featuring increased cardiac source TFs, resembling cluster 3 (Figures 6A-B, 6D-E, 7B and S7B-C); conversely, without WNTi, the mesoderm source TFs mirrored those present in cluster 5 (Figures 6C, 6F, 7C and S7A).

**FIGURE 7:**
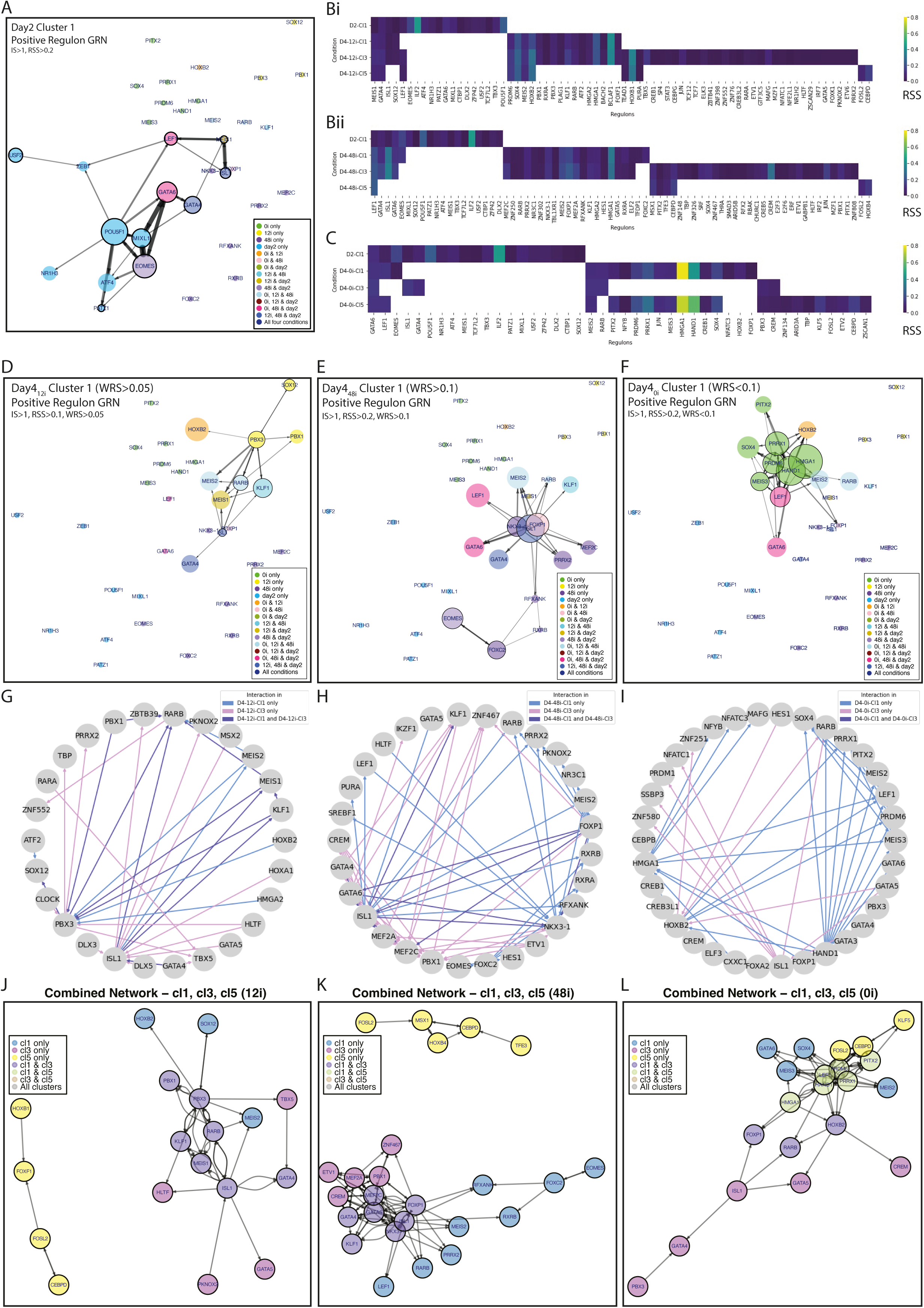
WNTi primes the transcriptional landscape of mesoderm (cluster 1) for cardiomyocyte differentiation. **A** Day 2 cluster 1 TF GRN showing positive regulon activity (node size). **B-C** Heatmaps of source regulons (RSS and IS filtered) in mesoderm (cluster 1, Cl1), cardiac progenitors (cluster 3, Cl3) and alternative cardiovascular cells (cluster 5, Cl5) at days (D) 2 and 4 under short (D4_12i_, **Bi**), (D4_48i_, **Bii**), or no (D4_0i_, **C**) WNTi. Colour code corresponds to regulon associated activity score. **D-F** Day 4 cluster 1 TF GRNs with WNTi-enabled positive regulon activity (node size) under Day4_12i_ (**D**), Day4_48i_ (**E**), and Day4_0i_ (**F**). **G-I** TF source-target interaction comparisons between Cl1 and Cl3 at day 4 under D4_12i_ (**G**), D4_48i_ (**H**), or D4_0i_ (**I**). Unique and shared interactions indicated. **J-L** Combined TF GRNs (positive regulon activity) for Cl1, Cl3 and Cl5 shown as a single network with cluster-specific colours (see legend; e.g. yellow = TFs present in Cl5 only, pink = TFs present exclusively in Cl3) for Day4_12i_ (**J**), Day4_48i_ (**K**), and Day4_0i_ (**L**). For panels **A, D-F**: colours indicate TF activity across all vs subset of conditions (see legend); a node border marks a source TF (no border = target only); interaction line thickness scales with IS.

To focus on WNTi effects, we imposed a WRS threshold (less stringent on D4_12i_). In keeping with the source regulon overlaps observed, WNTi induced a cardiac specification GRN within the mesoderm (Supplemental table 7 and Figures 6D-E and 7D-E), while a negative WRS in controls linked D4_0i_ mesoderm to cluster 5 (Fig. 7F and 6F). Intersecting mesoderm and cardiovascular networks identified shared source-target pairs (Figures 7G-I and S7D-F), revealing TFs that prime mesoderm for cardiomyocyte progenitor differentiation with WNTi versus those guiding alternative cardiovascular fates without WNTi. Short WNTi initiated a GRN centred around *PBX3* and *ISL1* (Figures 7D and 7G), reminiscent of the cardiomyocyte precursors’ GRN in the same condition. Longer WNTi triggered a second wave still centred on *ISL1* but where *FOXP1*, *NKX3*-1 played a strong role (Figures 7E and 7H). WNTi-treated mesoderm GRN showed little similarity to cluster 5 GRN (Figures S7D-E). Conversely, without WNTi, a GRN centred on *HMGA1*, *PRDM6* and *HAND1* emerged correlating with cluster 5 GRN but not with the cardiomyocyte progenitor cluster (3) (Figures 7F, 7I and S7F).

To visualise GRN assembly along the mesoderm-to-cardiomyocyte precursor trajectory, we merged cluster-level GRNs per sample (Supplemental Table 8, Figures 7J-L). Following WNTi (short or long), the cluster 5 GRN segregated, while clusters 1 and 3 interwove, consistent with WNTi priming mesoderm for a cardiomyocyte programme (Supplemental Table 8, Figures 7J-K). Without WNTi (D4_0i_), the mesoderm GRN connected with both clusters 3 and 5 GRN elements, indicating weak cell priming and broader lineage potential at day 4 (Figure 7L). Notably, *HOXB2*, the master regulon in D4_0i_ cluster 3 (Figures 6C and 6F), was a downstream positive target of cluster 5-specific TFs, supporting its potential repressing role in cardiomyogenesis (Figure 7L).

Overall, this analysis reveals WNTi patterns mesoderm to exit this progenitor state and commit directly to a cardiomyocyte fate by assembling a dedicated cardiogenic transcriptional programme. Without WNTi, mesoderm remains flexible and is not primed to any specific cardiovascular fate, with the resultant cardiomyocyte progenitors remaining entangled in competing GRNs, undermining their cardiogenic potential.

## DISCUSSION

WNT signalling is a key regulator of vertebrate heart development^15, 16^. Using a left ventricle-specific differentiation protocol^13^ we found that exogenous WNTi is dispensable for the emergence of some cardiovascular progenitor lineages, yet essential for efficient specification of cardiomyocyte precursors. Our findings demonstrate WNTi actively remodels the mesoderm transcriptional landscape to enable a sequential cardiomyocyte progenitor specification programme.

### Exogenous WNTi Is Required for Cardiac Specification In Vitro; Why are Endogenous WNT Signalling Inhibitors Not Sufficient?

Even without experimental WNTi, cells progress to cardiovascular progenitors, including some cardiomyocyte progenitors. In vivo, endoderm promotes cardiac differentiation by secreting WNT inhibitors, e.g. DKK1 downstream of a HHEX transcriptional programme^85^. As reported^86^, our cultures generate a small endodermal by-product expressing HHEX and DKK1, which may support initial progression to cardiovascular progenitors. These progenitors also express endogenous WNT inhibitors (*SFRP5*^42^ and *TMEM88*^43^) potentially sustaining initial differentiation towards cardiomyocyte lineages. However, this is insufficient and without experimental WNTi no robust cardiomyocyte progenitor specification and differentiation into beating cardiomyocytes takes place. Whether this reflects limited endodermal contribution in vitro remains unclear, and is a likely explanation given prior work shows hPSCs cardiomyocyte differentiation protocols benefit from co-culture with endoderm-like cells^87^. Future studies should define the tissue-context roles of endogenous WNT inhibitors.

### WNTi Directs Gene Expression for Specification and Differentiation of Cardiomyocytes; What’s the Alternative?

Lineage inference indicates that cardiovascular progenitors (clusters 2 and 7) represent a transitional state between mesoderm and two divergent fates: cardiomyocyte progenitors (cluster 3) or putative smooth muscle progenitors (cluster 5). This supports substantial plasticity in early mesoderm and these intermediates consistent with their mesenchymal, i.e. undefined, assignment by Capybara using gastrulating mouse embryo labels^29^. Fate choice depends on WNT context: experimental WNTi drives progression toward cardiomyocyte progenitors, reducing intermediates and alternative fates in line with reports that WNT-inhibited mesoderm favours cardiac identity^88^. Without WNTi, cells converge on cluster 5, which shows enrichment in smooth muscle terms, upregulation of the smooth muscle cell marker *ACTA2*^39^, and activation of a *PDRM16*-headed positive regulon (a smooth muscle TF^78^) —hallmarks of smooth-muscle development. Although Capybara did not label cluster 5 as smooth muscle, these features suggest it comprises of smooth muscle progenitors. Future work should probe the plasticity of these intermediates and their potential to adopt smooth muscle and other lineages without experimental WNTi.

### WNTi Initiates a Cardiomyocyte-promoting TF Programme in Mesoderm Cells

We inferred the GRN governing the mesoderm-to-cardiomyocyte-progenitor transition. In mesoderm, *EOMES* emerged as a key regulator, consistent with its role in cardiac mesoderm^89^, yet its target *MESP1* was not prominent, even after WNTi, despite its role in mesoderm specification and cardiac reprogramming^89, 90^. In contrast, TFs known in cardiomyocyte development, including *ISL1*^46^ and *MEIS2*^47, 48, 91^, were not only expressed but also acted as activators priming the mesoderm for cardiomyocyte differentiation. *GATA4*, a cardiac reprogramming factor^5, 8^, was also induced. Among other cardiac reprogramming factors^5, 8^, *MEF2C*, but not *TBX5,* was also activated in mesoderm after WNTi, as were *KLF1* and *NKX3-1*, which have not been linked previously to cardiomyocyte development. These factors later integrated into the cardiomyocyte progenitor GRN, likely reinforcing the cardiomyocyte progenitor programme and biasing mesoderm towards the cardiomyocyte lineage.

Besides evidence that WNTi initiates assembly of a cardiomyocyte specification GRN within the mesoderm, strong evidence for WNTi’s role in reshaping the mesodermal transcriptional landscape comes from the sustained downregulation and lack of connectivity of GRN elements associated with alternative cardiovascular lineages in WNTi-treated cultures. In contrast, without WNTi, the mesodermal GRN remains ambiguous, retaining connections to multiple cardiovascular programmes, suggesting cells keep lineage options open in the absence of a decisive signal. Future work should explore how WNTi-activated mesodermal genes instruct cardiomyocyte fate.

### WNTi Enables a Phased TF Network Establishment for Cardiomyocyte Specification

It was reassuring that our analysis identified TFs with well-established roles in heart development within the cardiomyocyte progenitor GRN, particularly after WNTi. Several (*GATA4*, *MEF2C*, *TBX5*) have been used for direct reprogramming of somatic cells into cardiomyocytes^5^. Notably, *GATA4* was an active regulator in the cardiomyocyte progenitor cluster irrespective of WNTi, even though cells without WNTi failed to differentiate and beat. Thus, while GATA4 is important for cardiogenic differentiation^50, 51^, it may not be central to the WNTi response requiring partners; the same may apply to *ISL1* and *MEIS2*. Unlike *GATA4*, *ISL1* is ineffective as a reprogramming factor^5, 8^, and is particularly important for secondary heart field structures^46^; thus, given our left ventricle-specific protocol^13^ and the first heart field origin of these cardiomyocytes^92^, *ISL1’s* limited pioneering role here is unsurprising without additional partners. Whether *MEIS2* promotes cardiac reprogramming remains unknown, although its paralogue *MEIS1* can facilitate it^93^.

By evaluating the WNTi-responsive epistasis and network hierarchy in the cardiomyocyte progenitor cluster, we further found that *ISL1* and *PBX3* may promote *GATA4* expression and initiate a cardiac GRN. However, without WNTi, they fail to establish a full GRN for subsequent differentiation. In contrast, with a short WNTi, *TBX5* becomes an activator, reinforcing *GATA4* expression and likely cooperating to promote cardiomyogenesis as previously reported^94^. Additionally, with WNTi *KLF1* also becomes an activator, and is reinforced by *ISL1* and *PBX3*, potentially opening cardiac genes’ chromatin given its pioneering role in other contexts^95^.

After longer WNTi, a mutually supporting GRN is established in cardiomyocyte progenitors, dominated by *MEF2C* (and *MEF2A*), *GATA6* (and *GATA4*), *ISL1*, *FOXP1* and *CREM*, and supported by *KLF1*, *NKX3*-*1* and *PBX1*. *KLF1* is a surprising WNTi-dependent regulator, potentially enabling assembly of the committed GRN, having previously only been linked to cardiogenesis in zebrafish heart-injury^68^. Similarly, *PBX1*^67^, *CREM*^70^, *NKX3-1*^96^ and *FOXP1*^97^, emerge as late WNTi responders, embedded in the committed GRN, despite limited prior links to cardiomyocyte specification. The presence of *FOXP1* was particularly unexpected given reports that *FOXP1* represses *NKX2*-5 and blocks cardiomyocyte proliferation in the developing mouse heart and in the border zone of injured mouse hearts^97, 98^. Whether this reflects interspecies differences remains to be determined.

Together, these data support a model in which without WNTi, only a partial cardiomyocyte GRN assembles (mainly *MEIS2*, *PBX3*, *ISL1* and *GATA4*) and is insufficient for cardiomyocyte differentiation. With WNTi, additional regulators are initially recruited (mainly *TBX5*, *KLF1* and *PBX1*) then others are added to establish a mutually supportive, committed GRN for cardiomyocyte differentiation (dominated by MEF2 genes, *GATA genes, FOXP1*, *ISL1* and *CREM*). Future work should confirm the role of these TFs in cardiomyocyte specification and test their required interactions for cardiomyogenesis.

### Why Does Absence of WNTi-treatment Preclude Differentiation of Cardiomyocyte Progenitors into Functional Cardiomyocytes?

A population of cardiac progenitors emerges even without exogenous WNTi, controlled by a GRN that shares TFs with the GRN of WNTi-derived cardiomyocyte progenitors, including *MEIS2*, *ISL1*, *PBX3* and *GATA4*. Additional TFs present in no-WNTi cardiomyocyte progenitors may compromise cardiomyocyte activators. *HOXB2* is a candidate because: it acts as a positive regulator across cardiovascular clusters with no or short WNTi but becomes a negative regulator within cardiomyocyte progenitors after prolonged WNTi, and its expression is markedly reduced following WNTi. Studies suggest HOXB2 can form a heterodimeric complex with PBX3 and MEIS genes^88^, potentially sequestering them from other partners and blocking progression of cardiomyocyte transcriptional programmes.

We also cannot exclude that without WNTi, activation of *GATA4*, *ISL1* and *PBX3* is insufficiently sustained for cardiomyocyte differentiation. Consistent, their expression was significantly higher in cardiomyocyte precursors after long WNTi than in controls, and *GATA4* is dosage-sensitive in cardiac morphogenesis^99^.

These mechanisms are not mutually exclusive and may together explain failed cardiomyocyte differentiation without WNTi. Future work should explore why WNTi-deprived cardiac progenitors do not progress to cardiomyocytes.

### WNTi Leads to Downregulation of TF Repressors of Cardiac Development

In addition to identifying transcriptional activators, we inferred transcriptional repressors. SCENIC detects repressors that are either expressed (targets repressed) or absent (targets de-repressed); we predominantly identified the latter, likely reflecting limits of transcript level detection, or RNA stability masking coordinated target downregulation.

We had anticipated confirming *MSX1* and CDX genes as negative regulators of cardiomyocyte differentiation^22, 23^. Although *MSX1*/MSX1 decreased with WNTi, *MSX1* was not identified as a repressor in cardiomyocyte precursors. Instead, increased *MSX1* expression and TF activator function in other cardiovascular clusters was observed, indicating that *MSX1* promotes alternative cardiovascular fates rather than directly repressing cardiomyocyte differentiation. This is consistent with *MSX1*-knockdown hPSCs-experiments showing that experimental WNTi is still required to produce cardiomyocytes^22^.

The principal repressor identified was HMGA1, whose reduced expression correlated with increased expression of its targets, the cardiomyocyte activators *MEF2C* and *GATA6*. This de-repression may be a result of cytoplasmatic shuttling of HMGA1 during cardiomyocyte-progenitor specification, as HMGA1 expression persisted irrespective of WNTi, while prolonged WNTi altered its subcellular localisation. This would be in keeping with reports of extra-nuclear HMGA1, and the view that dynamic subcellular localisation can shift its activity from chromatin remodelling toward roles in mRNA translation or signalling modulation^100^. Notably, *HMGA1* also acted as an activator in mesoderm and other cardiovascular lineages, suggesting that without WNTi it may restrict cardiomyocyte differentiation by maintaining mesoderm identity or alternative cardiovascular fates. Future work should confirm whether *HMGA1* acts as a repressive roadblock to cardiomyocyte differentiation and investigate its mechanism of action.

### Conclusion

We delineated a stepwise transcriptional programme for ventricular cardiomyocyte specification, identified new regulators including several activators and a few repressors, and mapped the TFs driving the mesoderm-to-cardiomyocyte-progenitor transition. Our findings show WNTi steers this trajectory by suppressing alternative lineage programmes and consolidating a cardiomyocyte-specific programme. Factors induced without WNTi fail to complete differentiation, marking them as peripheral or insufficient nodes in the specification GRN. These findings expose key control points in early cardiac fate decisions and provide a temporal framework to guide future dissection and engineering of cardiac lineage commitment.

## AUTHOR CONTRIBUTION STATEMENT

Conceptualization, S.H and A.S.B.; methodology, V.V. and A.S.B.; investigation, A.S.B., V.V., A.B., and E.F.; formal analysis, S.H., A.S.B., V.V., E.S, A.B, D.K. and F.S.; validation, V.V, A.B. and E.F.; visualization, A.S.B., V.V., A.B., E.S., D.K., and F.S.; data curation, E.S. F.S.; software, D.K. and F.S.; project administration, S.H. and A.S.B. writing – original draft, A.S.B. and S.H.; writing – review & editing, A.S.B., S.H., V.V., A.B., E.F, E.S., F.S.; funding acquisition, S.H. and A.S.B.; resources, A.S.B., and S.H.; supervision, A.S.B. and S.H..

## ACKNOWLEDGEMENTS

We thank Yvonne Turnbull (S.H. lab) for laboratory management; Dr Marie-Victoire Cosson (A.S.B. lab) for advice and support with hPSC culture; Elaina Collie-Duguid and Sophie Shaw (CGEBM) for advice and preliminary bioinformatics; Karishma Deggan for analysing TF expression across the published bulk RNA-seq time course^13^; and Genevia Technologies Oy (Tampere, Finland) for single-cell transcriptomics, statistical analyses, and data visualisations.

## REFERENCES

1. Tzahor E, Poss KD. Cardiac regeneration strategies: Staying young at heart. Science 2017;356:1035–1039.

2. NICE. What is the impact of CVD?, May 2023 ed: National Institute for Health and Care Excellence, 2023.

3. Sadek H, Olson EN. Toward the Goal of Human Heart Regeneration. Cell Stem Cell 2020;26:7–16.

4. Qian L, Huang Y, Spencer CI, Foley A, Vedantham V, Liu L, Conway SJ, Fu JD, Srivastava D. In vivo reprogramming of murine cardiac fibroblasts into induced cardiomyocytes. Nature 2012;485:593–598.

5. Ieda M, Fu JD, Delgado-Olguin P, Vedantham V, Hayashi Y, Bruneau BG, Srivastava D. Direct reprogramming of fibroblasts into functional cardiomyocytes by defined factors. Cell 2010;142:375–386.

6. Hausburg F, Jung JJ, Hoch M, Wolfien M, Yavari A, Rimmbach C, David R. (Re-)programming of subtype specific cardiomyocytes. Adv Drug Deliv Rev 2017;120:142–167.

7. Romero-Tejeda M, Fonoudi H, Weddle CJ, DeKeyser JM, Lenny B, Fetterman KA, Magdy T, Sapkota Y, Epting C, Burridge PW. A Novel Transcription Factor Combination for Direct Reprogramming to a Spontaneously Contracting Human Cardiomyocyte-like State. bioRxiv 2023.

8. Yamada Y, Sadahiro T, Ieda M. Development of direct cardiac reprogramming for clinical applications. J Mol Cell Cardiol 2023;178:1–8.

9. Murry CE, Keller G. Differentiation of embryonic stem cells to clinically relevant populations: lessons from embryonic development. Cell 2008;132:661–680.

10. Burridge PW, Matsa E, Shukla P, Lin ZC, Churko JM, Ebert AD, Lan F, Diecke S, Huber B, Mordwinkin NM, Plews JR, Abilez OJ, Cui B, Gold JD, Wu JC. Chemically defined generation of human cardiomyocytes. Nat Methods 2014;11:855–860.

11. Lee JH, Protze SI, Laksman Z, Backx PH, Keller GM. Human Pluripotent Stem Cell-Derived Atrial and Ventricular Cardiomyocytes Develop from Distinct Mesoderm Populations. Cell Stem Cell 2017;21:179–194 e174.

12. Lian X, Hsiao C, Wilson G, Zhu K, Hazeltine LB, Azarin SM, Raval KK, Zhang J, Kamp TJ, Palecek SP. Robust cardiomyocyte differentiation from human pluripotent stem cells via temporal modulation of canonical Wnt signaling. Proc Natl Acad Sci U S A 2012;109:E1848–1857.

13. Dark N, Cosson MV, Tsansizi LI, Owen TJ, Ferraro E, Francis AJ, Tsai S, Bouissou C, Weston A, Collinson L, Abi-Gerges N, Miller PE, MacLeod KT, Ehler E, Mitter R, Harding SE, Smith JC, Bernardo AS. Generation of left ventricle-like cardiomyocytes with improved structural, functional, and metabolic maturity from human pluripotent stem cells. Cell Rep Methods 2023;3:100456.

14. Yang D, Gomez-Garcia J, Funakoshi S, Tran T, Fernandes I, Bader GD, Laflamme MA, Keller GM. Modeling human multi-lineage heart field development with pluripotent stem cells. Cell Stem Cell 2022;29:1382–1401 e1388.

15. Hoppler S, Mazzotta S, Kuhl M. Wnt Signaling in Heart Organogenesis. Wnt Signaling in Development and Disease: Molecular Mechanisms and Biological Functions 2014:293–301.

16. Eisenberg LM, Eisenberg CA. Wnt signal transduction and the formation of the myocardium. Dev Biol 2006;293:305–315.

17. Yamaguchi TP, Takada S, Yoshikawa Y, Wu N, McMahon AP. T (Brachyury) is a direct target of Wnt3a during paraxial mesoderm specification. Genes Dev 1999;13:3185–3190.

18. Naito AT, Shiojima I, Akazawa H, Hidaka K, Morisaki T, Kikuchi A, Komuro I. Developmental stage-specific biphasic roles of Wnt/beta-catenin signaling in cardiomyogenesis and hematopoiesis. Proc Natl Acad Sci U S A 2006;103:19812–19817.

19. Ueno S, Weidinger G, Osugi T, Kohn AD, Golob JL, Pabon L, Reinecke H, Moon RT, Murry CE. Biphasic role for Wnt/beta-catenin signaling in cardiac specification in zebrafish and embryonic stem cells. Proc Natl Acad Sci U S A 2007;104:9685–9690.

20. Buikema JW, Mady AS, Mittal NV, Atmanli A, Caron L, Doevendans PA, Sluijter JP, Domian IJ. Wnt/beta-catenin signaling directs the regional expansion of first and second heart field-derived ventricular cardiomyocytes. Development 2013;140:4165–4176.

21. Mazzotta S, Neves C, Bonner RJ, Bernardo AS, Docherty K, Hoppler S. Distinctive Roles of Canonical and Noncanonical Wnt Signaling in Human Embryonic Cardiomyocyte Development. Stem Cell Reports 2016;7:764–776.

22. Rao J, Pfeiffer MJ, Frank S, Adachi K, Piccini I, Quaranta R, Arauzo-Bravo M, Schwarz J, Schade D, Leidel S, Scholer HR, Seebohm G, Greber B. Stepwise Clearance of Repressive Roadblocks Drives Cardiac Induction in Human ESCs. Cell Stem Cell 2016;18:341–353.

23. Munsterberg A, Hoppler S. WNT and BMP regulate roadblocks toward cardiomyocyte differentiation: lessons learned from embryos inform human stem cell differentiation. Stem Cell Investig 2016;3:33.

24. Zheng GX, Terry JM, Belgrader P, Ryvkin P, Bent ZW, Wilson R, Ziraldo SB, Wheeler TD, McDermott GP, Zhu J, Gregory MT, Shuga J, Montesclaros L, Underwood JG, Masquelier DA, Nishimura SY, Schnall-Levin M, Wyatt PW, Hindson CM, Bharadwaj R, Wong A, Ness KD, Beppu LW, Deeg HJ, McFarland C, Loeb KR, Valente WJ, Ericson NG, Stevens EA, Radich JP, Mikkelsen TS, Hindson BJ, Bielas JH. Massively parallel digital transcriptional profiling of single cells. Nat Commun 2017;8:14049.

25. Stuart T, Butler A, Hoffman P, Hafemeister C, Papalexi E, Mauck WM, 3rd, Hao Y, Stoeckius M, Smibert P, Satija R. Comprehensive Integration of Single-Cell Data. Cell 2019;177:1888–1902 e1821.

26. Conway JR, Lex A, Gehlenborg N. UpSetR: an R package for the visualization of intersecting sets and their properties. Bioinformatics 2017;33:2938–2940.

27. Cao J, Spielmann M, Qiu X, Huang X, Ibrahim DM, Hill AJ, Zhang F, Mundlos S, Christiansen L, Steemers FJ, Trapnell C, Shendure J. The single-cell transcriptional landscape of mammalian organogenesis. Nature 2019;566:496–502.

28. Yu G, Wang LG, Han Y, He QY. clusterProfiler: an R package for comparing biological themes among gene clusters. OMICS 2012;16:284–287.

29. Pijuan-Sala B, Griffiths JA, Guibentif C, Hiscock TW, Jawaid W, Calero-Nieto FJ, Mulas C, Ibarra-Soria X, Tyser RCV, Ho DLL, Reik W, Srinivas S, Simons BD, Nichols J, Marioni JC, Gottgens B. A single-cell molecular map of mouse gastrulation and early organogenesis. Nature 2019;566:490–495.

30. Durinck S, Spellman PT, Birney E, Huber W. Mapping identifiers for the integration of genomic datasets with the R/Bioconductor package biomaRt. Nat Protoc 2009;4:1184–1191.

31. Kong W, Fu YC, Holloway EM, Garipler G, Yang X, Mazzoni EO, Morris SA. Capybara: A computational tool to measure cell identity and fate transitions. Cell Stem Cell 2022;29:635–649 e611.

32. Aibar S, Gonzalez-Blas CB, Moerman T, Huynh-Thu VA, Imrichova H, Hulselmans G, Rambow F, Marine JC, Geurts P, Aerts J, van den Oord J, Atak ZK, Wouters J, Aerts S. SCENIC: single-cell regulatory network inference and clustering. Nat Methods 2017;14:1083–1086.

33. Kumar N, Mishra B, Athar M, Mukhtar S. Inference of Gene Regulatory Network from Single-Cell Transcriptomic Data Using pySCENIC. Methods Mol Biol 2021;2328:171–182.

34. Csárdi G, Nepusz T, Traag V, Horvát S, Zanini F, Noom D, Müller K. igraph: Network Analysis and Visualization in R., version 1.5.1 ed. https://igraph.org, 2025:R package.

35. Hunter JD. Matplotlib: A 2D graphics environment. Comput Sci Eng 2007;9:90–95.

36. Aric A. Hagberg DASaPJS. Exploring network structure, dynamics, and function using NetworkX. Python in Science Conference 2008;Proceeding of the 7th Python in Science Conference (SciPy2008):11–15.

37. Waskom ML. seaborn: statistical data visualization. The JOurnal of Open SOurce Software 2021;6:3021.

38. Cheung C, Bernardo AS, Trotter MW, Pedersen RA, Sinha S. Generation of human vascular smooth muscle subtypes provides insight into embryological origin-dependent disease susceptibility. Nat Biotechnol 2012;30:165–173.

39. Guo DC, Papke CL, Tran-Fadulu V, Regalado ES, Avidan N, Johnson RJ, Kim DH, Pannu H, Willing MC, Sparks E, Pyeritz RE, Singh MN, Dalman RL, Grotta JC, Marian AJ, Boerwinkle EA, Frazier LQ, LeMaire SA, Coselli JS, Estrera AL, Safi HJ, Veeraraghavan S, Muzny DM, Wheeler DA, Willerson JT, Yu RK, Shete SS, Scherer SE, Raman CS, Buja LM, Milewicz DM. Mutations in smooth muscle alpha-actin (ACTA2) cause coronary artery disease, stroke, and Moyamoya disease, along with thoracic aortic disease. Am J Hum Genet 2009;84:617–627.

40. Dalton S. Linking the Cell Cycle to Cell Fate Decisions. Trends Cell Biol 2015;25:592–600.

41. La Manno G, Soldatov R, Zeisel A, Braun E, Hochgerner H, Petukhov V, Lidschreiber K, Kastriti ME, Lonnerberg P, Furlan A, Fan J, Borm LE, Liu Z, van Bruggen D, Guo J, He X, Barker R, Sundstrom E, Castelo-Branco G, Cramer P, Adameyko I, Linnarsson S, Kharchenko PV. RNA velocity of single cells. Nature 2018;560:494–498.

42. Fujii M, Sakaguchi A, Kamata R, Nagao M, Kikuchi Y, Evans SM, Yoshizumi M, Shimono A, Saga Y, Kokubo H. Sfrp5 identifies murine cardiac progenitors for all myocardial structures except for the right ventricle. Nat Commun 2017;8:14664.

43. Palpant NJ, Pabon L, Rabinowitz JS, Hadland BK, Stoick-Cooper CL, Paige SL, Bernstein ID, Moon RT, Murry CE. Transmembrane protein 88: a Wnt regulatory protein that specifies cardiomyocyte development. Development 2013;140:3799–3808.

44. Xian L, Georgess D, Huso T, Cope L, Belton A, Chang YT, Kuang W, Gu Q, Zhang X, Senger S, Fasano A, Huso DL, Ewald AJ, Resar LMS. HMGA1 amplifies Wnt signalling and expands the intestinal stem cell compartment and Paneth cell niche. Nat Commun 2017;8:15008.

45. Arrington CB, Dowse BR, Bleyl SB, Bowles NE. Non-synonymous variants in pre-B cell leukemia homeobox (PBX) genes are associated with congenital heart defects. Eur J Med Genet 2012;55:235–237.

46. Cai CL, Liang X, Shi Y, Chu PH, Pfaff SL, Chen J, Evans S. Isl1 identifies a cardiac progenitor population that proliferates prior to differentiation and contributes a majority of cells to the heart. Dev Cell 2003;5:877–889.

47. Paige SL, Thomas S, Stoick-Cooper CL, Wang H, Maves L, Sandstrom R, Pabon L, Reinecke H, Pratt G, Keller G, Moon RT, Stamatoyannopoulos J, Murry CE. A temporal chromatin signature in human embryonic stem cells identifies regulators of cardiac development. Cell 2012;151:221–232.

48. Shanmugam K, Green NC, Rambaldi I, Saragovi HU, Featherstone MS. PBX and MEIS as non-DNA-binding partners in trimeric complexes with HOX proteins. Mol Cell Biol 1999;19:7577–7588.

49. Cao C, Li L, Zhang Q, Li H, Wang Z, Wang A, Liu J. Nkx2.5: a crucial regulator of cardiac development, regeneration and diseases. Front Cardiovasc Med 2023;10:1270951.

50. Oka T, Maillet M, Watt AJ, Schwartz RJ, Aronow BJ, Duncan SA, Molkentin JD. Cardiac-specific deletion of Gata4 reveals its requirement for hypertrophy, compensation, and myocyte viability. Circ Res 2006;98:837–845.

51. Pikkarainen S, Tokola H, Kerkela R, Ruskoaho H. GATA transcription factors in the developing and adult heart. Cardiovasc Res 2004;63:196–207.

52. Gingold JA, Fidalgo M, Guallar D, Lau Z, Sun Z, Zhou H, Faiola F, Huang X, Lee DF, Waghray A, Schaniel C, Felsenfeld DP, Lemischka IR, Wang J. A genome-wide RNAi screen identifies opposing functions of Snai1 and Snai2 on the Nanog dependency in reprogramming. Mol Cell 2014;56:140–152.

53. Paul S, Zhang X, He JQ. Homeobox gene Meis1 modulates cardiovascular regeneration. Semin Cell Dev Biol 2020;100:52–61.

54. Niwa H, Miyazaki J, Smith AG. Quantitative expression of Oct-3/4 defines differentiation, dedifferentiation or self-renewal of ES cells. Nat Genet 2000;24:372–376.

55. Wu SP, Cheng CM, Lanz RB, Wang T, Respress JL, Ather S, Chen W, Tsai SJ, Wehrens XH, Tsai MJ, Tsai SY. Atrial identity is determined by a COUP-TFII regulatory network. Dev Cell 2013;25:417–426.

56. Jiang Y, Zhang Z. OVOL2: an epithelial lineage determiner with emerging roles in energy homeostasis. Trends Cell Biol 2023;33:824–833.

57. Castano J, Aranda S, Bueno C, Calero-Nieto FJ, Mejia-Ramirez E, Mosquera JL, Blanco E, Wang X, Prieto C, Zabaleta L, Mereu E, Rovira M, Jimenez-Delgado S, Matson DR, Heyn H, Bresnick EH, Gottgens B, Di Croce L, Menendez P, Raya A, Giorgetti A. GATA2 Promotes Hematopoietic Development and Represses Cardiac Differentiation of Human Mesoderm. Stem Cell Reports 2019;13:515–529.

58. Duong TB, Holowiecki A, Waxman JS. Retinoic acid signaling restricts the size of the first heart field within the anterior lateral plate mesoderm. Dev Biol 2021;473:119–129.

59. Liu F, van den Broek O, Destree O, Hoppler S. Distinct roles for Xenopus Tcf/Lef genes in mediating specific responses to Wnt/beta-catenin signalling in mesoderm development. Development 2005;132:5375–5385.

60. Sorrell MR, Dohn TE, D’Aniello E, Waxman JS. Tcf7l1 proteins cell autonomously restrict cardiomyocyte and promote endothelial specification in zebrafish. Dev Biol 2013;380:199–210.

61. Li Y, Du J, Deng S, Liu B, Jing X, Yan Y, Liu Y, Wang J, Zhou X, She Q. The molecular mechanisms of cardiac development and related diseases. Signal Transduct Target Ther 2024;9:368.

62. Takahashi K, Tanabe K, Ohnuki M, Narita M, Ichisaka T, Tomoda K, Yamanaka S. Induction of pluripotent stem cells from adult human fibroblasts by defined factors. Cell 2007;131:861–872.

63. Narayan S, Bryant G, Shah S, Berrozpe G, Ptashne M. OCT4 and SOX2 Work as Transcriptional Activators in Reprogramming Human Fibroblasts. Cell Rep 2017;20:1585–1596.

64. Stellato M, Dewenter M, Rudnik M, Hukara A, Ozsoy C, Renoux F, Pachera E, Gantenbein F, Seebeck P, Uhtjaerv S, Osto E, Razansky D, Klingel K, Henes J, Distler O, Blyszczuk P, Kania G. The AP-1 transcription factor Fosl-2 drives cardiac fibrosis and arrhythmias under immunofibrotic conditions. Commun Biol 2023;6:161.

65. Biasin V, Marsh LM, Egemnazarov B, Wilhelm J, Ghanim B, Klepetko W, Wygrecka M, Olschewski H, Eferl R, Olschewski A, Kwapiszewska G. Meprin beta, a novel mediator of vascular remodelling underlying pulmonary hypertension. J Pathol 2014;233:7–17.

66. Huang GN, Thatcher JE, McAnally J, Kong Y, Qi X, Tan W, DiMaio JM, Amatruda JF, Gerard RD, Hill JA, Bassel-Duby R, Olson EN. C/EBP transcription factors mediate epicardial activation during heart development and injury. Science 2012;338:1599–1603.

67. Chang CP, Stankunas K, Shang C, Kao SC, Twu KY, Cleary ML. Pbx1 functions in distinct regulatory networks to pattern the great arteries and cardiac outflow tract. Development 2008;135:3577–3586.

68. Ogawa M, Geng FS, Humphreys DT, Kristianto E, Sheng DZ, Hui SP, Zhang Y, Sugimoto K, Nakayama M, Zheng D, Hesselson D, Hodson MP, Bogdanovic O, Kikuchi K. Kruppel-like factor 1 is a core cardiomyogenic trigger in zebrafish. Science 2021;372:201–205.

69. Pon JR, Marra MA. MEF2 transcription factors: developmental regulators and emerging cancer genes. Monotargeted 2016;7:2297–2312.

70. Schulte JS, Fehrmann E, Tekook MA, Kranick D, Fels B, Li N, Wehrens XH, Heinick A, Seidl MD, Schmitz W, Muller FU. Cardiac expression of the CREM repressor isoform CREM-IbDeltaC-X in mice leads to arrhythmogenic alterations in ventricular cardiomyocytes. Basic Res Cardiol 2016;111:15.

71. Doppler SA, Werner A, Barz M, Lahm H, Deutsch MA, Dressen M, Schiemann M, Voss B, Gregoire S, Kuppusamy R, Wu SM, Lange R, Krane M. Myeloid zinc finger 1 (Mzf1) differentially modulates murine cardiogenesis by interacting with an Nkx2.5 cardiac enhancer. PLoS One 2014;9:e113775.

72. Shekhar A, Lin X, Lin B, Liu FY, Zhang J, Khodadadi-Jamayran A, Tsirigos A, Bu L, Fishman GI, Park DS. ETV1 activates a rapid conduction transcriptional program in rodent and human cardiomyocytes. Sci Rep 2018;8:9944.

73. Hovanes K, Li TW, Munguia JE, Truong T, Milovanovic T, Lawrence Marsh J, Holcombe RF, Waterman ML. Beta-catenin-sensitive isoforms of lymphoid enhancer factor-1 are selectively expressed in colon cancer. Nat Genet 2001;28:53–57.

74. Davidson EH. The regulatory genome : gene regulatory networks in development and evolution. Burlington, MA ; San Diego: Academic, 2006.

75. Moerman T, Aibar Santos S, Bravo Gonzalez-Blas C, Simm J, Moreau Y, Aerts J, Aerts S. GRNBoost2 and Arboreto: efficient and scalable inference of gene regulatory networks. Bioinformatics 2019;35:2159–2161.

76. Balamurugan K, Sterneck E. The many faces of C/EBPdelta and their relevance for inflammation and cancer. Int J Biol Sci 2013;9:917–933.

77. Zhang S, Li P, Li J, Gao J, Qi Q, Dong G, Liu X, Jiao Q, Wang Y, Du L, Zhan H, Xu S, Wang C. Chromatin accessibility uncovers KRAS-driven FOSL2 promoting pancreatic ductal adenocarcinoma progression through up-regulation of CCL28. Br J Cancer 2023;129:426–443.

78. Davis CA, Haberland M, Arnold MA, Sutherland LB, McDonald OG, Richardson JA, Childs G, Harris S, Owens GK, Olson EN. PRISM/PRDM6, a transcriptional repressor that promotes the proliferative gene program in smooth muscle cells. Mol Cell Biol 2006;26:2626–2636.

79. McFadden DG, Barbosa AC, Richardson JA, Schneider MD, Srivastava D, Olson EN. The Hand1 and Hand2 transcription factors regulate expansion of the embryonic cardiac ventricles in a gene dosage-dependent manner. Development 2005;132:189–201.

80. Campione M, Ros MA, Icardo JM, Piedra E, Christoffels VM, Schweickert A, Blum M, Franco D, Moorman AF. Pitx2 expression defines a left cardiac lineage of cells: evidence for atrial and ventricular molecular isomerism in the iv/iv mice. Dev Biol 2001;231:252–264.

81. Cserjesi P, Lilly B, Bryson L, Wang Y, Sassoon DA, Olson EN. MHox: a mesodermally restricted homeodomain protein that binds an essential site in the muscle creatine kinase enhancer. Development 1992;115:1087–1101.

82. Ryan K, Garrett N, Mitchell A, Gurdon JB. Eomesodermin, a key early gene in Xenopus mesoderm differentiation. Cell 1996;87:989–1000.

83. Ng ES, Azzola L, Sourris K, Robb L, Stanley EG, Elefanty AG. The primitive streak gene Mixl1 is required for efficient haematopoiesis and BMP4-induced ventral mesoderm patterning in differentiating ES cells. Development 2005;132:873–884.

84. Zeineddine D, Papadimou E, Chebli K, Gineste M, Liu J, Grey C, Thurig S, Behfar A, Wallace VA, Skerjanc IS, Puceat M. Oct-3/4 dose dependently regulates specification of embryonic stem cells toward a cardiac lineage and early heart development. Dev Cell 2006;11:535–546.

85. Foley AC, Mercola M. Heart induction by Wnt antagonists depends on the homeodomain transcription factor Hex. Genes Dev 2005;19:387–396.

86. Zhao M, Tang Y, Zhou Y, Zhang J. Deciphering Role of Wnt Signalling in Cardiac Mesoderm and Cardiomyocyte Differentiation from Human iPSCs: Four-dimensional control of Wnt pathway for hiPSC-CMs differentiation. Sci Rep 2019;9:19389.

87. Mummery C, Ward-van Oostwaard D, Doevendans P, Spijker R, van den Brink S, Hassink R, van der Heyden M, Opthof T, Pera M, de la Riviere AB, Passier R, Tertoolen L. Differentiation of human embryonic stem cells to cardiomyocytes: role of coculture with visceral endoderm-like cells. Circulation 2003;107:2733–2740.

88. den Hartogh SC, Wolstencroft K, Mummery CL, Passier R. A comprehensive gene expression analysis at sequential stages of in vitro cardiac differentiation from isolated MESP1-expressing-mesoderm progenitors. Sci Rep 2016;6:19386.

89. Costello I, Pimeisl IM, Drager S, Bikoff EK, Robertson EJ, Arnold SJ. The T-box transcription factor Eomesodermin acts upstream of Mesp1 to specify cardiac mesoderm during mouse gastrulation. Nat Cell Biol 2011;13:1084– 1091.

90. Chan SS, Shi X, Toyama A, Arpke RW, Dandapat A, Iacovino M, Kang J, Le G, Hagen HR, Garry DJ, Kyba M. Mesp1 patterns mesoderm into cardiac, hematopoietic, or skeletal myogenic progenitors in a context-dependent manner. Cell Stem Cell 2013;12:587–601.

91. Stankunas K, Shang C, Twu KY, Kao SC, Jenkins NA, Copeland NG, Sanyal M, Selleri L, Cleary ML, Chang CP. Pbx/Meis deficiencies demonstrate multigenetic origins of congenital heart disease. Circ Res 2008;103:702–709.

92. Kelly RG, Buckingham ME, Moorman AF. Heart fields and cardiac morphogenesis. Cold Spring Harb Perspect Med 2014;4.

93. Wang J, Jiang X, Zhao L, Zuo S, Chen X, Zhang L, Lin Z, Zhao X, Qin Y, Zhou X, Yu XY. Lineage reprogramming of fibroblasts into induced cardiac progenitor cells by CRISPR/Cas9-based transcriptional activators. Acta Pharm Sin B 2020;10:313–326.

94. Ang YS, Rivas RN, Ribeiro AJS, Srivas R, Rivera J, Stone NR, Pratt K, Mohamed TMA, Fu JD, Spencer CI, Tippens ND, Li M, Narasimha A, Radzinsky E, Moon-Grady AJ, Yu H, Pruitt BL, Snyder MP, Srivastava D. Disease Model of GATA4 Mutation Reveals Transcription Factor Cooperativity in Human Cardiogenesis. Cell 2016;167:1734–1749 e1722.

95. Magor GW, Gillinder KR, Huang S, Ilsley MD, Bell C, Perkins AC. KLF1 Acts As a Pioneer Transcription Factor Via SMARCA4 to Open Chromatin and Facilitate Redeployment of an Enhancer Complex Containing GATA1 and SCL. Blood 2022;140.

96. Schneider A, Brand T, Zweigerdt R, Arnold H. Targeted disruption of the Nkx3.1 gene in mice results in morphogenetic defects of minor salivary glands: parallels to glandular duct morphogenesis in prostate. Mech Dev 2000;95:163–174.

97. Zhang Y, Li S, Yuan L, Tian Y, Weidenfeld J, Yang J, Liu F, Chokas AL, Morrisey EE. Foxp1 coordinates cardiomyocyte proliferation through both cell-autonomous and nonautonomous mechanisms. Genes Dev 2010;24:1746– 1757.

98. Wang Y, Wang X, Fang J, Chen X, Xu T, Zhuang T, Peng S, Bao W, Wu W, Lu Y, Wang H, Tomlinson B, Chan P, Zhuang S, Zhang Q, Zhang L, Liu Z, Pi J, Zhang Y, Liu J. Cardiomyocyte Foxp1-Specific Deletion Promotes Post-injury Heart Regeneration via Targeting Usp20-HIF1a-Hand1 Signaling Pathway. Adv Sci (Weinh) 2025;12:e2412124.

99. Pu WT, Ishiwata T, Juraszek AL, Ma Q, Izumo S. GATA4 is a dosage-sensitive regulator of cardiac morphogenesis. Dev Biol 2004;275:235–244.

100. Pujals M, Resar L, Villanueva J. HMGA1, Moonlighting Protein Function, and Cellular Real Estate: Location, Location, Location! Biomolecules 2021;11.

