## Supplemental Figures and Methods for "WNT inhibition primes the transcriptional landscape of mesoderm to initiate a phased ventricular cardiomyocyte specification programme"

**SUPPLEMENTAL MATERIAL**

SUPPLEMENTAL FIGURES LEGENDS:

**FIGURE S1: Inhibition of WNT signalling directs differentiation towards cardiomyocyte progenitors and away from alternative cardiovascular fates**

**A** Flow cytometry analysis of TNNT2 expression in day 12 samples grown in the presence of short (D3.5–D4, 12i) or long (D2–D4, 48i) WNTi, or in its absence (no WNTi control). **B** UMAP charts depicting cell cycle state of mesoderm cells (day 2, D2), or day 4 samples grown in the presence of short ($D4_{12i}$) or long ($D4_{48i}$) WNTi, or in its absence ($D4_{0i}$). **C** UMAP charts of marker genes prominent in different clusters, including: pluripotency-associated genes (**Ci**), mesoderm-associated genes (**Cii**), endoderm-associated genes (**Ciii**), and neuro-mesodermal associated genes (**Civ**). **D** Upset plot of relationship groupings among clusters based on shared marker gene expression. **E** UMAP chart (of combined samples) illustrating Capybara analysis using annotated cell fates proposed for early mouse embryogenesis[32]. Note cardiomyocyte label predominantly in highlighted area of cluter 3. **F** UMAP charts depicting cardiomyocytes as labelled using Capybara based on early mouse embryogenesis[32] in all day 4 samples including controls ($D4_{0i}$) and those grown in the presence of WNT inhibitior ($D4_{12i}$ and $D4_{48i}$). **G-H** UMAP charts of cluster 5 (**G**) and cluster 3 (**H**) marker genes in day 4 samples grown in the presence of short ($D4_{12i}$) or long ($D4_{48i}$) WNTi, or in its absence ($D4_{0i}$).

**FIGURE S2: WNTi Promotes Direct Cardiomyocyte Progenitor Differentiation and Structural Gene Expression**

**A** Violin plots of WNTi-linked gene expression within cluster 3 in day 2 mesoderm samples and in day 4 samples grown in the presence of short ($Day4_{12i}$) or long ($Day4_{48i}$) WNTi, or in its absence ($Day4_{0i}$). **B** Validation of WNTi-linked gene expression by real time RT-PCR in cultures differentiated without WNTi (0i), shown as a time course analysis. One-way ANOVA performed on a n=3. Data presented as mean +/- standard deviation. **C** GO terms associated with cluster 3 after short WNTi (i.e., $D4_{12i}$). **D** NES plots from GSEA analysis of cell proliferation associated terms decreased in cluster 3.

**Figure S3: WNTi Modifies the Transcriptional Landscape of Cardiac Progenitor Cells**

**A** Heatmap showing time course analysis of TF genes differentially expressed in cluster 3 upon WNTi. Gene expression was plotted against a bulk RNA-sequencing dataset of differentiating left ventricle cardiomyocytes[13]. **B** UMAP charts of *MSX1 CDX2* and *CDX1* expression in day 2 mesoderm samples and in day 4 samples grown in the presence of short ($Day4_{12i}$) or long ($Day4_{48i}$) WNTi, or in its absence ($Day4_{0i}$). **C-D** Immunofluorescence staining showing MSX1 and HMGA1 (**C**) or LUM and ISL1 (**D**) expression in day 4 cultures grown with short WNTi ($D4_{12i}$). Cell nuclei were counterstained with DAPI. **E** Graphs quantifying ISL1 expression based on immunistaining. Quantification was determined based on percentage of ISL1 expressing cells (**Ei**) and ISL1 fluorescence intensity (**Eii**). Welch's ANOVA performed on a n=2 with each having 2 biological replicates. Data presented as mean +/- standard deviation.

**FIGURE S4: WNTi Restricts Expression of Repressor TFs, Revealing Novel Inhibitors of Cardiomyocyte Differentiation.**

**A-B** Heatmap of negative regulon activity in day (D) 4 control ($D4_{0i}$) samples grown in the absence of WNTi (**A**) or in the presence of long WNTi ($D4_{48i}$) (**B**) in the indicated clusters. **C** Comparison between TF expression (UMAP chart, left) and regulon activity (ridge plot, right) of: *SNAI2* with SNAI2(-) (**Ci**); *MEIS1* with MEIS1(-) (**Cii**); and *GATA4* with GATA4(-) (**Ciii**). **D** Bubble plot of top negative regulons in cluster 3 in day 4 samples grown with ($D4_{12i}$, $D4_{48i}$) and without ($D4_{0i}$) WNTi. Warmer colours indicate cluster-specificity of regulon activity, the thickness of the outline indicates regulon activity, and the size of the bubble indicates the number of potential downstream regulated genes in the dataset. E Comparison between *TCF7L1* and *TCF7L2* expression (top) and their negative regulon activity (bottom) across clusters.

**FIGURE S5: WNTi Activates Expression of TFs Promoting Cardiac Specification, Revealing Novel Drivers of Cardiomyocyte Differentiation**

**A-B** Heatmap of positive regulon activity in day 4 samples cultured without ($D4_{0i}$, **A**) or with long WNTi ($D4_{48i}$, **B**); clusters analysed are indicated. **C-D** Regulon activity ridge plot of POU5F1(+) (**C**) and SOX2(+) (**D**). **E** UMAP comparison of *CEBPD* gene expression with CEBPD(+) regulon activity and the expression of *HAND1*, one of the inferred target genes for this regulon. **F** UMAP comparison of *FOSL2* gene expression with FOSL2(+) regulon activity and the expression of *ADAM19*, one of the inferred target genes for this regulon. **G** UMAP comparison of *MEIS2* gene expression (**Gi**) with MEIS2(+) regulon activity (**Gii**). **H** Time-course of *PBX3* (**Hi**), ISL1 (**Hii**) and MEF2C (**Hiii**) expression measured by real-time RT-PCR. One-way ANOVA with data presented as mean +/- standard deviation; n=3. **I** Comparison of TF expression (as indicated) in clusters (violin plots, top) with corresponding positive regulon activity (UMAPs, bottom) following standard WNTi ($D4_{48i}$).

**FIGURE S6: WNTi Establishes a TF Gene Regulatory Network for promoting Cardiomyocyte Cluster 3 versus the alternative fate in cluster 5**

**A-B** TF GRN associated with transcriptional repressors linked to negative regulon activity (size of node) in cluster 3 focusing on day 4 samples grown in the absence ($Day4_{0i}$, **A**) or the presence of long WNTi ($Day4_{48i}$, **B**). Colour coding indicates whether a TF is active across all or only a subset of experimental samples (see legend). A border around the TF bubble denotes a source TF, while absence of a border indicates it functions only as a target under the defined thresholds. Line thickness reflects the importance score of each source–target interaction. **C-D** TF GRN associated with WNTi-repressed positive regulon activity (size of node) in cluster 5, focussing on day 4 samples grown in the presence of short ($Day4_{12i}$)or long ($Day4_{48i}$) WNTi. Colour code, bubble boarder and line thickness were defined as above.

**FIGURE S7: WNTi primes the transcriptional landscape of mesoderm (cluster 1) for cardiomyocyte differentiation**

**A-C** TF GRN associated with positive regulon activity (size of node) in cluster 1 of day 4 samples grown in the presence of short ($Day4_{12i}$, **A**), long ($Day4_{48i}$, **B**) or absence ($Day4_{0i}$, **C**) of WNTi. Colour code, bubble boarder and line thickness were defined as above. Colour coding indicates whether a TF is active across all or only a subset of experimental samples (see legend). A border around the TF bubble denotes a source TF, while absence of a border indicates it functions only as a target under the defined thresholds. Line thickness reflects the importance score of each source–target interaction. **D-F** Comparison of TF source-target interaction between the mesoderm cluster (Cl1) and the alternative cardiovascular fate cluster (Cl5) in day 4 samples grown in the presence of short ($D4_{12i}$, **D**) or long ($D4_{48i}$, **E**) WNTi, or in its absence ($D4_{0i}$, **F**). Unique and shared TF interactions are indicated.

SUPPLEMENTARY TABLES:

**Suppl. Table 1:** List of genes enriched in each cluster

**Suppl. Table 2:** Gene set enrichment analysis per cluster

**Suppl. Table 3:** Differential gene expression analysis in cluster 3

**Suppl. Table 4:** List of regulons per cluster, including potential target genes

**Suppl. Table 5:** GRN feed table for cluster 3 including RSS, activity score and WNT response score

**Suppl. Table 6:** GRN feed table for cluster 5 including RSS, activity score and WNT response score

**Suppl. Table 7:** GRN feed table for cluster 1 including RSS, activity score and WNT response score

**Suppl. Table 8:** GRN combined summary table with filtered TFs driving GRNs in clusters per sample type.

#### MATERIALS AND METHODS

##### Human pluripotent stem cell lines: Origin, characterization and maintenance

PluriTest (Thermo Fisher Scientific) and KaryoStat were used to assess pluripotency and genetic stability respectively prior to cryopreservation. In addition, routine pluripotency testing was performed using BD Stemflow™ Human Pluripotent Stem Cell Analysis Kit (BD Bioscience), along with low pass sequencing karyotype analysis, and standard immunostaining for pluripotency markers OCT3/4, SOX2 and NANOG was conducted. Karyotype stability of these lines was further monitored through more frequent low pass sequencing.

##### Differentiation of human pluripotent stem cell lines to cardiomyocytes

Once cells reached roughly 70-80% confluency, RPMI 1640 media was added (ThermoFisher) supplemented with B27 minus insulin (ThermoFisher), Activin A (5 ng/ml, R&D), CHIR999021 (2 μM, Sellek Chemikals), BMP4 (5-6 ng/ml, R&D) and FGF2 (5 ng/ml, R&D) for 24 hours and replaced with RPMI 1640 media supplemented with B27 minus insulin for one additional day. On Day 2, media was replaced with RPMI 1640 media supplemented with B27 minus insulin, L-ascorbic acid (65 μg/ml, Sigma), and the Wnt inhibitor IWR1 (1 μM, Sigma) or IWP2 (5 μM, Sigma) for 48 hours (also known as our 48i treated samples). WNTi studies required variation of WNTi period (see results). From Day 6, media was replaced every two days with RPMI 1640 supplemented with B27 with insulin (ThermoFisher) and L-ascorbic Acid.

**Immunocytochemistry**
Cells were fixed using 4% paraformaldehyde (ThermoFisher Scientific) for 10 minutes at room temperature and permeabilised with 0.1% Triton X-100 (Sigma) for 10 minutes and then blocked using 5% Normal Donkey Serum (Sigma) in PBS (Sigma) for 1 hour at room temperature. Fixed cells were incubated with primary antibodies (see below), diluted in 5% donkey serum (Sigma) and incubated overnight at 4 °C in a humidity chamber. This was followed by three PBS washes and adding the fluorochrome conjugated secondary antibodies (see below), diluted in 5% Donkey Serum in PBS for 1h at room temperature. Cells were washed three times with PBS and DAPI (1:10,000; ThermoFisher Scientific) added, incubated for 10 minutes and washed twice. Images were acquired using a Zeiss Confocal microscope (Zeiss) and Zeiss Zen software and analysed using ImageJ software.

**Immunostaining Primary Antibody Table**

| **Primary Antibody** | **Species** | **Dilution** | **Source** | **Identifier** |
|---|---|---|---|---|
| **AACTN** | Mouse | 1:200 | Merck | A7811 |
| **ISL1** | Goat | 1:100 | R&D Systems | AF1837 |
| **TNNT2** | Rabbit | 1:200 | Abcam | AB45932 |
| **TNNT2** | Mouse | 1:200 | Invitrogen | MA5-12960 |
| **NKX2.5** | Goat | 1:75 | Santa Cruz Biotechnology | Sc-8697 |
| **ISL1** | Mouse | 1:75 | Abcam | AB86472 |
| **GATA4** | Rabbit | 1:150 | Cell Signalling Technology | 36966 |
| **HMGA1** | Rabbit | 1:100 | Abcam | AB129153 |
| **LUM** | Rabbit | 1:200 | Invitrogen | MA5-29402 |
| **EOMES** | Rabbit | 1:100 | Cell Signalling Technology | 81493 |
| **COL3A1** | Rabbit | 1:200 | NOVUS | NB600-594 |
| **POU5F1** | Mouse | 1:200 | Santa Cruz | SC-5279 |
| **NANOG** | Goat | 1:200 | R&D | AF1997 |
| **SOX2** | Rabbit | 1:200 | Millipore | AB5603 |
| **MSX1** | Goat | 1:100 | R&D Systems | AF5045 |

**Immunostaining Secondary Antibody Table**

| Secondary Antibodies | Species | Dilution | Source | Identifier |
|---|---|---|---|---|
| Anti-goat Alexa 488 | Donkey | 1:500 | ThermoFisher | A-11055 |
| Anti-mouse Alexa 488 | Donkey | 1:500 | ThermoFisher | A-21202 |
| Anti-rabbit Alexa 488 | Donkey | 1:500 | ThermoFisher | A-21206 |
| Anti-goat Alexa 594 | Donkey | 1:500 | ThermoFisher | A-11058 |
| Anti-mouse Alexa 594 | Donkey | 1:500 | ThermoFisher | A-21203 |

| | | | | |
|---|---|---|---|---|
| Anti-rabbit Alexa 594 | Donkey | 1:500 | ThermoFisher | A-21207 |

**Flow cytometry**

Samples were fixed and stained as described in main manuscript. Pluripotent assessments using POU5F1, NANGO, SOX2, SSEA1 and SSEA4 were done regularly but data not included in manuscript. Assessment of cardiomyocyte differentiation efficiency is reported.

**Flow Cytometry Antibody Table**

| Primary Antibodies | Species | Dilution | Source | Identifier |
|---|---|---|---|---|
| POU5F1-PecCP5.5 (FC)<br>Isotype-PerCP5.5 | Mouse | 20 µl / 1x$10^6$ cells | BD-Bioscience | 560794 |
| NANOG-647 (FC)<br>Isotype-647 (FC) | Mouse | 5 µl / 1x$10^6$ cells | BD-Bioscience | 561300 |
| SOX2-488 (FC)<br>Isotype-488 (FC) | Mouse | 2.5 µl / 1x$10^6$ cells | BD-Bioscience | 561593 |
| SSEA1-PE (FC)<br>Isotype-PE (FC) | Mouse | 20 µl / 1x$10^6$ cells | BD-Bioscience | 560886 |
| SSEA4-647 (FC)<br>Isotype-647 (FC) | Mouse | 5 µl / 1x$10^6$ cells | BD-Bioscience | 563119 |
| TNNT2-PE (FC)<br>Isotype-PE(FC) | Mouse | 1:200 / 1x$10^6$ cells | BD-Bioscience | 564767<br>554680 |

**RNA extraction, cDNA synthesis and RT-pPCR**

RNA was quantified (Nanodrop) and 1 µg of RNA used for cDNA synthesis using the Maxima First Strand cDNA synthesis kit (Thermo Fisher Scientific) according to manufacturer's instructions and using dsDNase treatment. The cDNA samples were then diluted (1:100) in DNase/RNase-free water and 2 µl of diluted cDNA used per reaction by real-time qPCR using GoTaq Master Mix (Promega) as per manufacturer's instructions, including qPCR cycling conditions. Primers (see Primer Table below) were checked and accepted if efficiency was 100% ±10%. Analyses were carried out with an internal standard curve per plate and using PBGD1 as a reference gene.

**Primer table**

| Gene | Primer Sequence (5' to 3') | Product Size (bp) | RefSeq Accession Number |
|---|---|---|---|
| *ISL1* | F: TGCTTTTCAGCAACTGGTCA<br>R: TGAATGTTCCTCATGCCTCA | 143 | NM_002202.2 |
| *MEF2C* | F: ACGAGGATTATGGATGAACG<br>R: TGGCATACTGGAACAGCTTG | 151 | NM_002397.4 |
| *MEIS1* | F: CGGTGGCCACACGTCACACA<br>R: TCGTCACCTGTGCTGGGGGA | 90 | NM_002398.3 |
| *PBGD1* (housekeeper) | F: ATTACCCCGGGAGACTGAAC<br>R: GGCTGTTGCTTGGACTTCTC | 130 | NM_000190.3 |

| | | | |
|---|---|---|---|
| *PBX3* | F: CCCTCCGTCATGTTATCAATCAG<br>R: GCCAGCCTCCATTAGCATTTA | 99 | NM_006195.6 |
| *SFRP5* | F: CCGCTGGGACAAGAAGAATAA<br>R: CCCGTAGAAGAAAGGGTAGTAGA | 91 | NM_003015.3 |
| *TMEM88* | F: TGGCTGCCTTCAATCTTCTC<br>R: ACTGAGTGGCAGAGGAA | 97 | NM_203411.2 |

**Single Cell RNA sequencing**
Sample viability was between 84 and 87% and mean cell concentrations were optimal for loading between 880 and 1200 cells per µl. For each sample approximately 6000 cells were loaded for a targeted recovery of 3000 cells. Samples were loaded onto individual lanes of a Chip B and run on a Chromium Controller (10x Genomics). Libraries were then prepared and indexed according to manufacturer's instructions (Chromium Single Cell 3′ Reagent Kit, v3 Chemistry, 10x Genomics). Libraries were pooled equally by initial estimated cell count and sequenced on a NextSeq 500 using High Output v2.5 (150 cycle) reagents and flow cell. CellRanger[98] was used to perform demultiplexing, barcode processing and single-cell 3′ gene counts, resulting in a recovery of between ~2700 and 4100 cells per sample (~50 – 65%).

**Sample integration and clustering**
Feature barcode matrices for each sample were imported into Seurat v3 and cells with fewer than 200 or more than 7500 transcribed genes were removed. Cells with more than 25% mitochondrial reads were also removed. Expression for each gene was log normalized and variable features were computed (these are genes which display a high degree of variation within the dataset) prior to further processing using Seurat[99] defaults. Data integration was performed using the CCA method. Dimensionality reduction (PCA, UMAP) and clustering were performed accordingly followed by graph-based clustering (Louvain method), for multiple resolutions, with 0.5 having the best match according to expected cell types.

**Capybara label transferring**
Using our humanized dataset, we constructed high-resolution references at the cell-type and sub-type levels (construct.high.res.reference), computed background and test quadratic-programming identity scores (single.round.QP.analysis), and obtained empirical p-values via randomization (percentage.calc). Scores were binarized (binarization.mann.whitney) and converted to per-cell calls (binary.to.classification). Multi-identity cells were curated (multi.id.curate.qp), and transition scores were calculated from the curated multi-ID set (transition.score). For per-class fraction scoring, we generated pseudo-bulk references with Seurat's AverageExpression and re-scored the query. The resulting Capybara fractions and calls were written back to the Seurat object metadata for downstream analyses.

**Regulon analysis with SCENIC**
In order to validate WNT-regulated TFs we used SCENIC software[45,46] to identify groups of TF-co-regulated genes within our single-cell transcriptomics dataset. SCENIC infers a regulon, named after the TF, comprising predicted direct targets identified by co-expression and motif enrichment. Regulons are split by sign:

TF (+), positive, lists targets whose expression increases with the TF (putative activation), whereas TF (-), negative, lists targets whose expression decreases as the TF increases (putative repression, i.e., targets that are higher when the TF is low). Naturally, a single TF may have both (+) and (-) regulons in different cellular contexts, capturing distinct sets of activated and repressed genes.

This workflow generated regulon specificity scores (RSS) to quantify how selectively each regulon characterizes individual clusters, with higher RSS indicating stronger cluster-specific regulatory influence. Per-cell regulon activity was computed with AUCell, which tests whether a regulon's targets are enriched among the top-expressed genes in each cell. Cluster-level activity was then obtained by aggregating per-cell AUCell scores within each cluster, providing a quantitative measure of regulon activation across clusters.

**Inference of Gene Regulatory Networks (GRNs)**

SCENIC-defined TF regulons were analysed to infer higher-order TF–TF relationships by adding an edge whenever a TF's regulon contained another TF as a predicted target These interactions can be unidirectional or reciprocal, activating or repressing. From the full set of inferred TF gene interactions we retained those with higher importance scores and strong cluster-specificity, as indicated by regulon specificity scores (RSS) in the clusters of interest.

For "igraph" library[105,106] generated GRNs, input data was used to define the GRN look with: node size defined by log2(activity score *100 +1); thickness of line guiding interactions defined by log2(importance +1) / log2(global_max_importance + 1) * 5; and arrow head size defined by line_thickness / 4 . Global_max_importance was defined as the highest importance score in a certain cluster. Only source nodes displayed a line around them. Targets were kept in the GRN if they were fed by more than one source in case of transcriptional activator GRNs but all targets were kept if GRNs displayed transcriptional repressors.

For Matplotlib[36] generated GRNs, input data was used to define GRN look with: node size based on RSS score, and thickness of the lines based on importance score.

For NetworkX generated GRNs, input data was used to define the GRN look with: line colour was defined based on the intersection (or absence of thereof) between clusters.

Seaborn[38] was used to plot network comparisons based on sources and their activity score per sample and cluster. Sources were filtered for RSS and importance score.

All filters applied to data are specified in the GRN filter table, any exceptions are stated in the figures.

**GRN filter table**

| | Source/ Target Sign | Source_ RSS | Importance Score | Target_RS S | Source_ WRS_ Z.score* | Target_ WRS_ Z.score* |
|---|---|---|---|---|---|---|
| D2-Cl1 | + | ≥0.2 | ≥1 | ≥0.2 | | |
| $D4_{0i}$-Cl1 | + | ≥0.1 | ≥1 | ≥0.1 | ≤-0.1 | |
| $D4_{12i}$-Cl1 | + | ≥0.1 | ≥1 | ≥0.1 | ≥0.05 | |
| $D4_{48i}$-Cl1 | + | ≥0.2 | ≥1 | ≥0.1 | ≥0.1 | |
| $D4_{0i}$-Cl3 | + | ≥0.1 | ≥1 | | ≥0.1 | ≥0 |
| $D4_{12i}$-Cl3 | + | ≥0.1 | ≥1 | | ≥0.1 | ≥0 |
| $D4_{48i}$-Cl3 | + | ≥0.2 | ≥1 | ≥0.2 | ≥0.1 | ≥0.1 |
| $D4_{0i}$-Cl5 | + | ≥0.1 | ≥1 | ≥0.1 | ≤-0.05 | ≤-0.05 |
| $D4_{12i}$-Cl5 | + | ≥0.1 | ≥1 | ≥0.1 | ≤-0.05 | ≤-0.05 |
| $D4_{48i}$-Cl5 | + | ≥0.1 | ≥1 | ≥0.1 | ≤-0.05 | ≤-0.05 |

*Not all figures used WRS, this is indicated in the figure/figure legend.

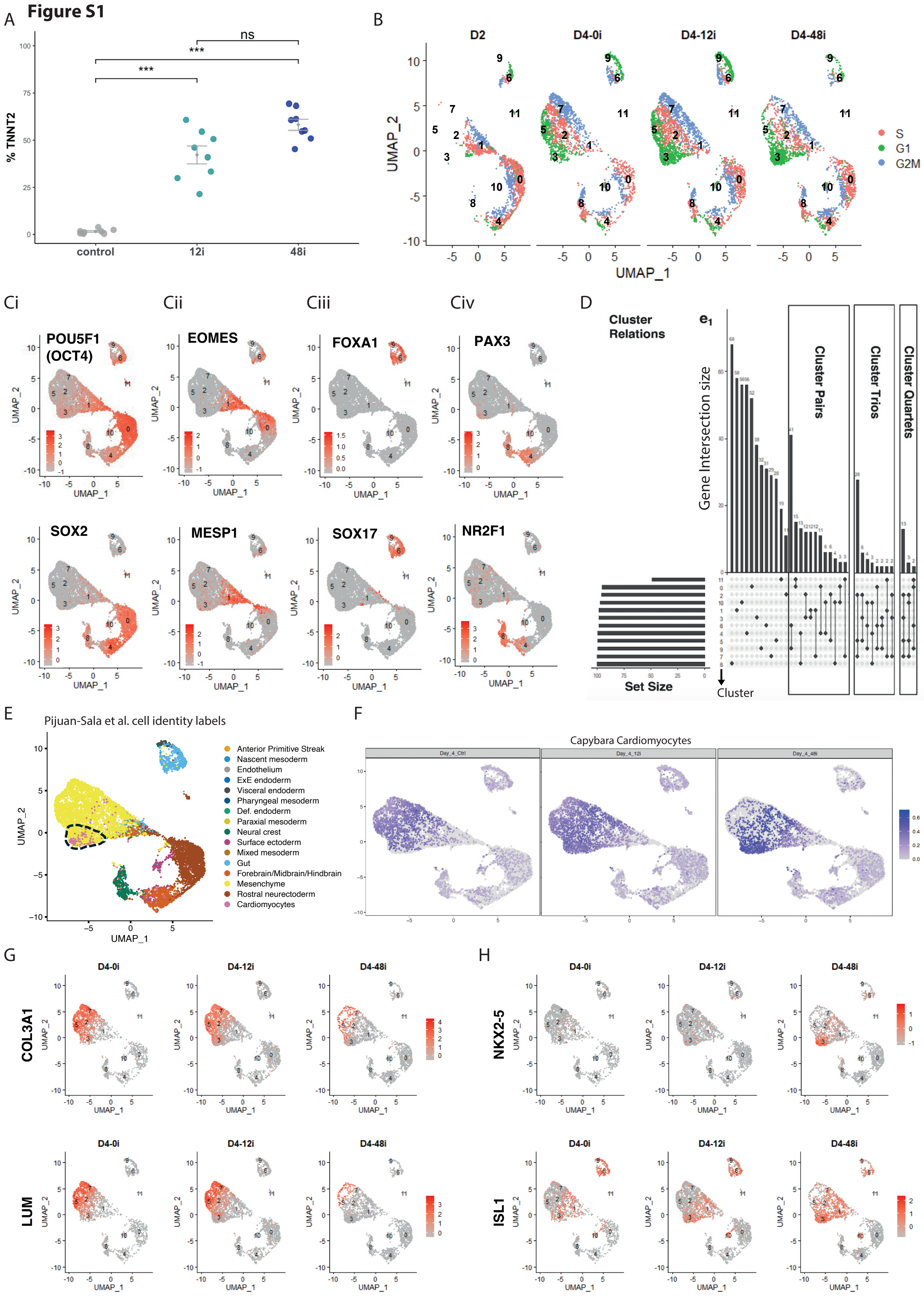
Figure S1
A
% TNNT2
ns
***
***
control
12i
48i
B
D2
D4-0i
D4-12i
D4-48i
UMAP_2
UMAP_1
S
G1
G2M
Ci
Cii
Ciii
Civ
POU5F1 (OCT4)
EOMES
FOXA1
PAX3
SOX2
MESP1
SOX17
NR2F1
D
Cluster Relations
Gene Intersection size
Cluster Pairs
Cluster Trios
Cluster Quartets
Set Size
Cluster
E
Pijuan-Sala et al. cell identity labels
Anterior Primitive Streak
Nascent mesoderm
Endothelium
ExE endoderm
Visceral endoderm
Pharyngeal mesoderm
Def. endoderm
Paraxial mesoderm
Neural crest
Surface ectoderm
Mixed mesoderm
Gut
Forebrain/Midbrain/Hindbrain
Mesenchyme
Rostral neurectoderm
Cardiomyocytes
F
Capybara Cardiomyocytes
Day_4_Ctrl
Day_4_12i
Day_4_48i
G
COL3A1
LUM
H
NKX2-5
ISL1

#### Figure S2

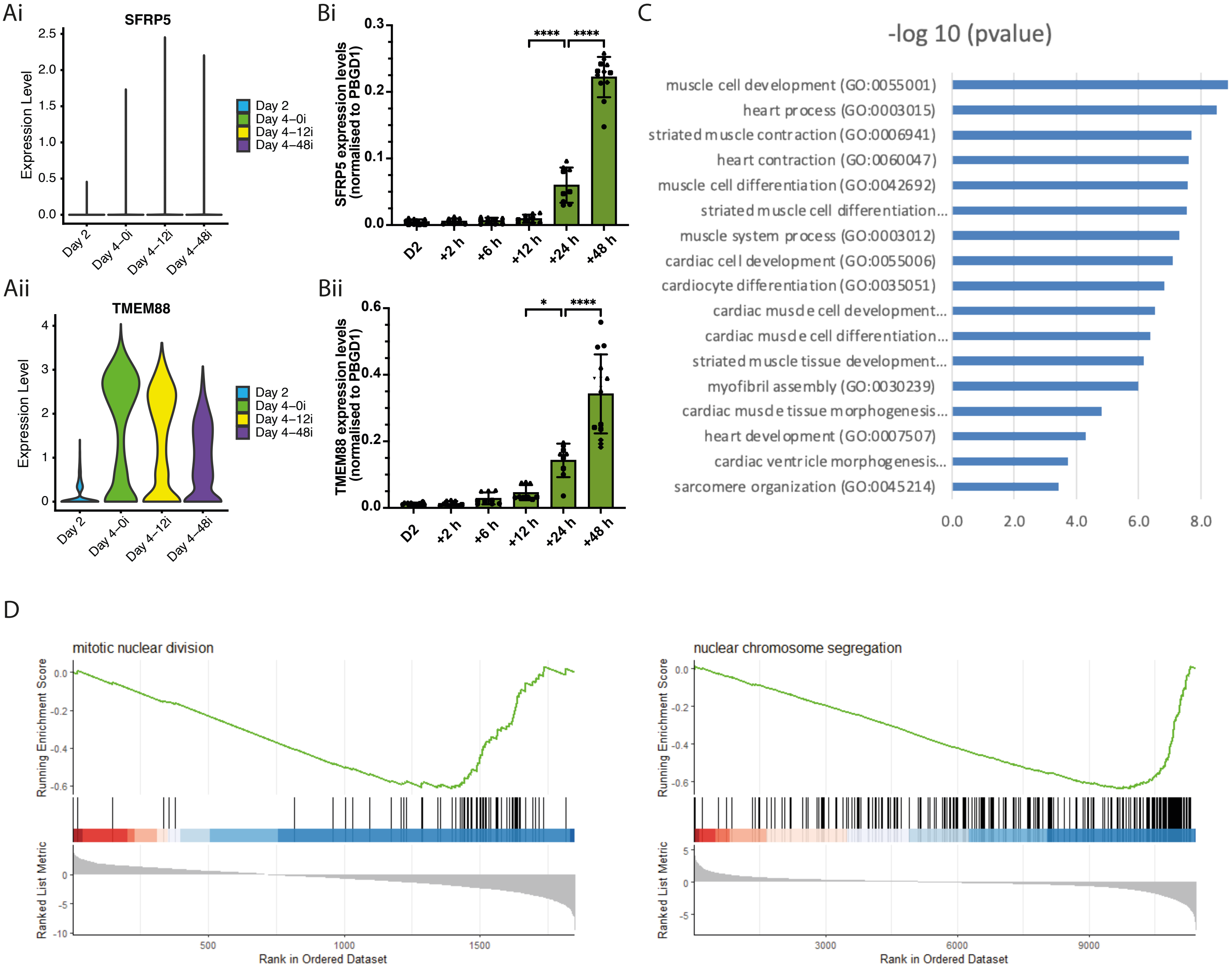

Ai
SFRP5
Expression Level
2.5
2.0
1.5
1.0
0.5
0.0
Day 2
Day 4–0i
Day 4–12i
Day 4–48i
Bi
SFRP5 expression levels (normalised to PBGD1)
0.3
0.2
0.1
0.0
****
****
D2
+2 h
+6 h
+12 h
+24 h
+48 h
Aii
TMEM88
Expression Level
4
3
2
1
0
Day 2
Day 4–0i
Day 4–12i
Day 4–48i
Bii
TMEM88 expression levels (normalised to PBGD1)
0.6
0.4
0.2
0.0
*
****
D2
+2 h
+6 h
+12 h
+24 h
+48 h
C
-log 10 (pvalue)
muscle cell development (GO:0055001)
heart process (GO:0003015)
striated muscle contraction (GO:0006941)
heart contraction (GO:0060047)
muscle cell differentiation (GO:0042692)
striated muscle cell differentiation...
muscle system process (GO:0003012)
cardiac cell development (GO:0055006)
cardiocyte differentiation (GO:0035051)
cardiac muscle cell development...
cardiac muscle cell differentiation...
striated muscle tissue development...
myofibril assembly (GO:0030239)
cardiac muscle tissue morphogenesis...
heart development (GO:0007507)
cardiac ventricle morphogenesis...
sarcomere organization (GO:0045214)
0.0
2.0
4.0
6.0
8.0
D
mitotic nuclear division
Running Enrichment Score
Ranked List Metric
Rank in Ordered Dataset
500
1000
1500
nuclear chromosome segregation
Running Enrichment Score
Ranked List Metric
Rank in Ordered Dataset
3000
6000
9000

#### Figure S3

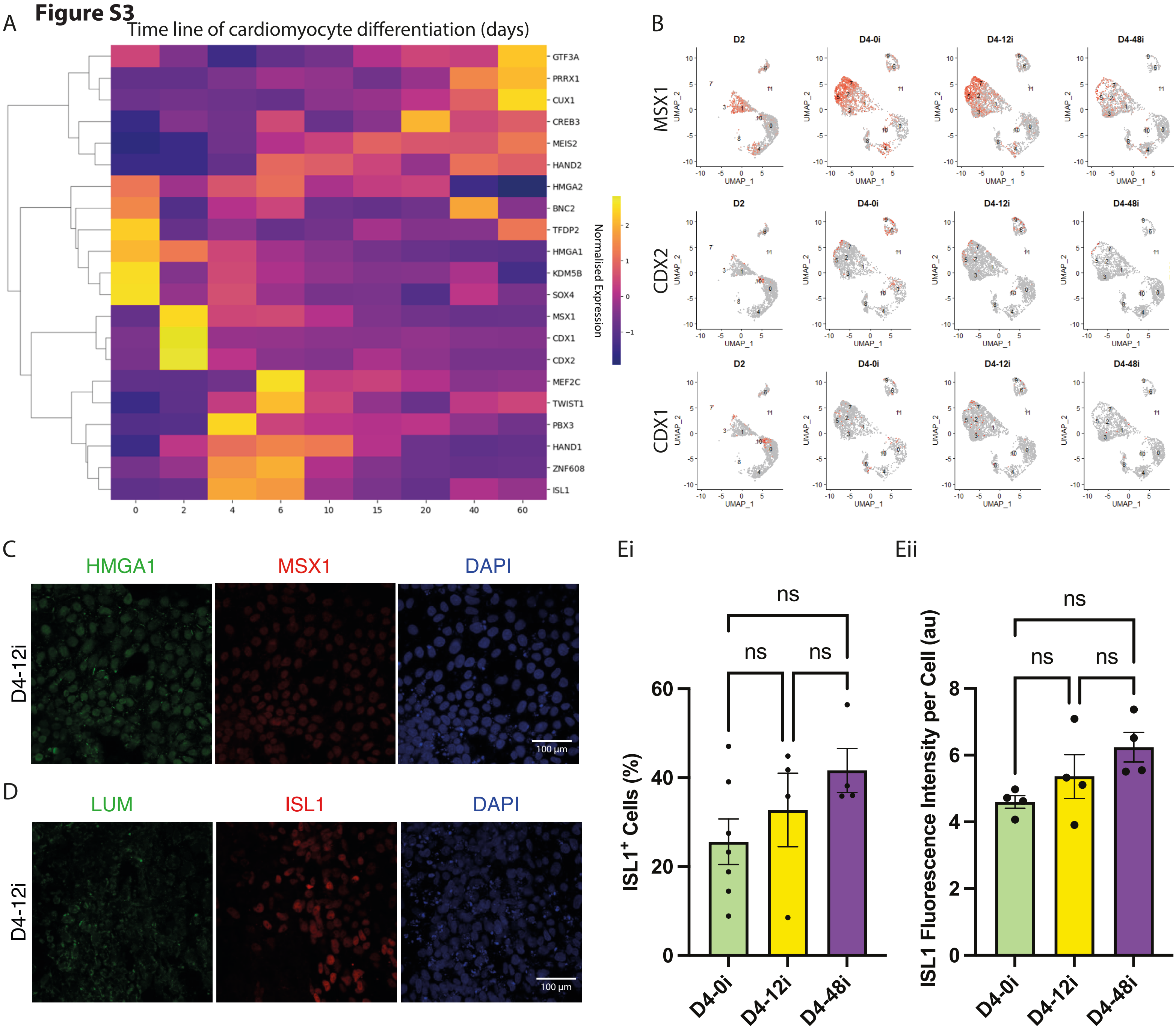
A
Time line of cardiomyocyte differentiation (days)
GTF3A
PRRX1
CUX1
CREB3
MEIS2
HAND2
HMGA2
BNC2
TFDP2
HMGA1
KDM5B
SOX4
MSX1
CDX1
CDX2
MEF2C
TWIST1
PBX3
HAND1
ZNF608
ISL1
Normalised Expression
B
D2
D4-0i
D4-12i
D4-48i
MSX1
CDX2
CDX1
UMAP_1
UMAP_2
C
HMGA1
MSX1
DAPI
D4-12i
100 µm
D
LUM
ISL1
DAPI
D4-12i
100 µm
Ei
Eii
ns
ISL1+ Cells (%)
ISL1 Fluorescence Intensity per Cell (au)
D4-0i
D4-12i
D4-48i

### Figure S4

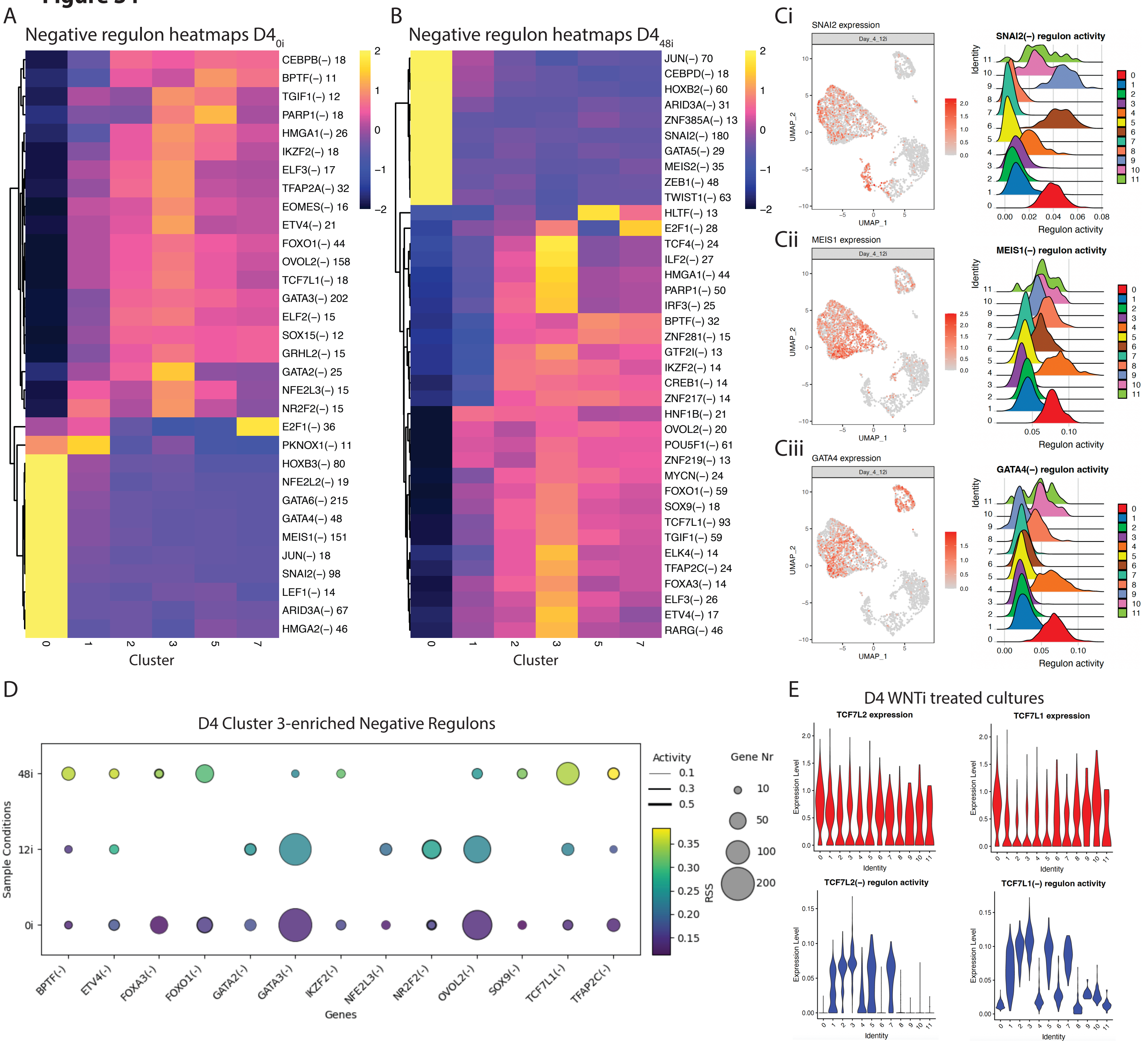
A
Negative regulon heatmaps D4 0i
CEBPB(–) 18
BPTF(–) 11
TGIF1(–) 12
PARP1(–) 18
HMGA1(–) 26
IKZF2(–) 18
ELF3(–) 17
TFAP2A(–) 32
EOMES(–) 16
ETV4(–) 21
FOXO1(–) 44
OVOL2(–) 158
TCF7L1(–) 18
GATA3(–) 202
ELF2(–) 15
SOX15(–) 12
GRHL2(–) 15
GATA2(–) 25
NFE2L3(–) 15
NR2F2(–) 15
E2F1(–) 36
PKNOX1(–) 11
HOXB3(–) 80
NFE2L2(–) 19
GATA6(–) 215
GATA4(–) 48
MEIS1(–) 151
JUN(–) 18
SNAI2(–) 98
LEF1(–) 14
ARID3A(–) 67
HMGA2(–) 46
Cluster
B
Negative regulon heatmaps D4 48i
JUN(–) 70
CEBPD(–) 18
HOXB2(–) 60
ARID3A(–) 31
ZNF385A(–) 13
SNAI2(–) 180
GATA5(–) 29
MEIS2(–) 35
ZEB1(–) 48
TWIST1(–) 63
HLTF(–) 13
E2F1(–) 28
TCF4(–) 24
ILF2(–) 27
HMGA1(–) 44
PARP1(–) 50
IRF3(–) 25
BPTF(–) 32
ZNF281(–) 15
GTF2I(–) 13
IKZF2(–) 14
CREB1(–) 14
ZNF217(–) 14
HNF1B(–) 21
OVOL2(–) 20
POU5F1(–) 61
ZNF219(–) 13
MYCN(–) 24
FOXO1(–) 59
SOX9(–) 18
TCF7L1(–) 93
TGIF1(–) 59
ELK4(–) 14
TFAP2C(–) 24
FOXA3(–) 14
ELF3(–) 26
ETV4(–) 17
RARG(–) 46
Cluster
Ci
SNAI2 expression
Day_4_12i
SNAI2(–) regulon activity
Cii
MEIS1 expression
MEIS1(–) regulon activity
Ciii
GATA4 expression
GATA4(–) regulon activity
UMAP_1
UMAP_2
Identity
Regulon activity
D
D4 Cluster 3-enriched Negative Regulons
Activity
Gene Nr
RSS
Sample Conditions
48i
12i
0i
BPTF(-)
ETV4(-)
FOXA3(-)
FOXO1(-)
GATA2(-)
GATA3(-)
IKZF2(-)
NFE2L3(-)
NR2F2(-)
OVOL2(-)
SOX9(-)
TCF7L1(-)
TFAP2C(-)
Genes
E
D4 WNTi treated cultures
TCF7L2 expression
TCF7L1 expression
TCF7L2(–) regulon activity
TCF7L1(–) regulon activity
Expression Level
Identity

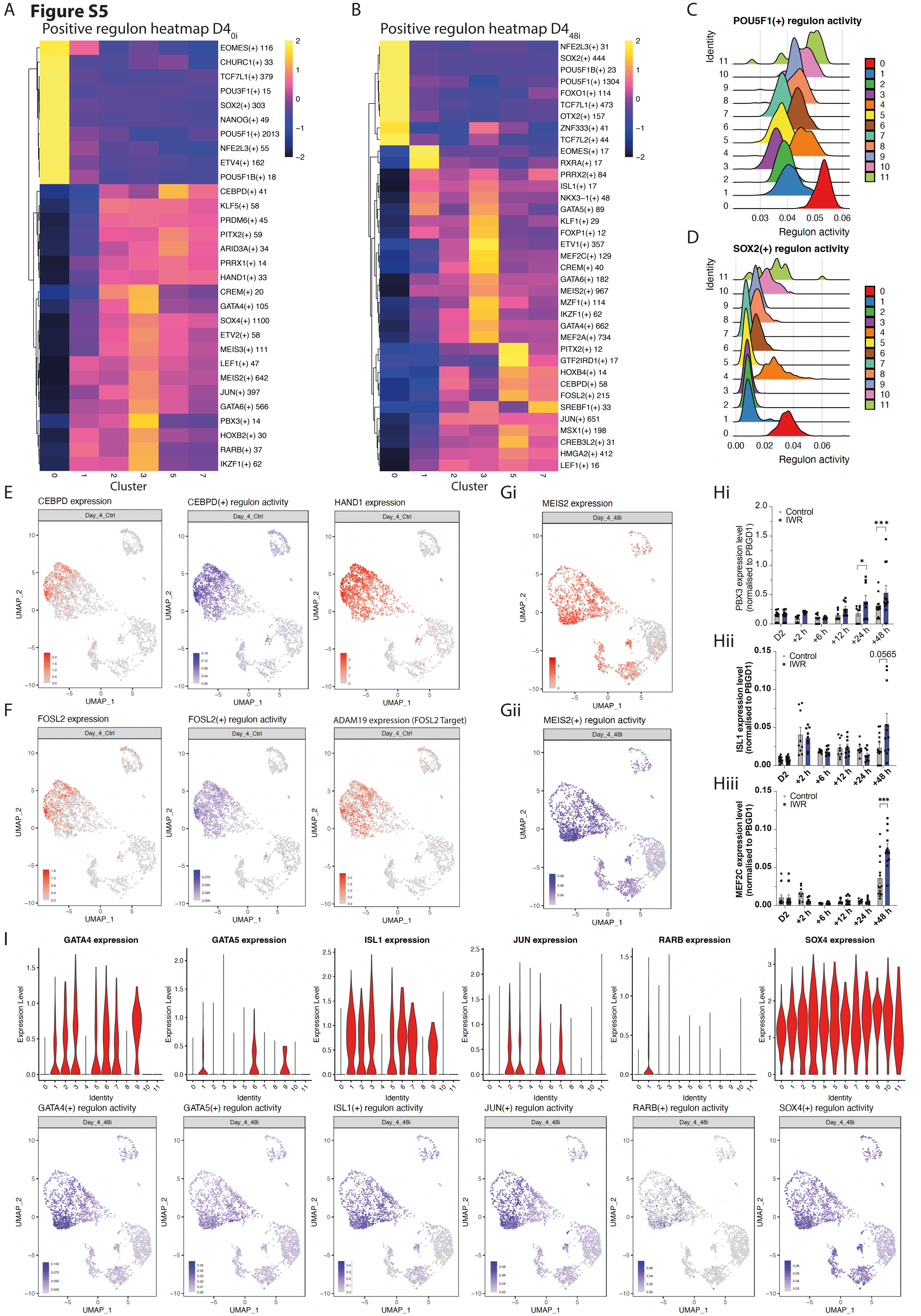
Figure S5
A
Positive regulon heatmap D4_0i
B
Positive regulon heatmap D4_48i
C
POU5F1(+) regulon activity
D
SOX2(+) regulon activity
Identity
Regulon activity
Cluster
E
CEBPD expression
CEBPD(+) regulon activity
HAND1 expression
F
FOSL2 expression
FOSL2(+) regulon activity
ADAM19 expression (FOSL2 Target)
Gi
MEIS2 expression
Gii
MEIS2(+) regulon activity
Day_4_Ctrl
Day_4_48i
UMAP_1
UMAP_2
Hi
PBX3 expression level (normalised to PBGD1)
Hii
ISL1 expression level (normalised to PBGD1)
Hiii
MEF2C expression level (normalised to PBGD1)
Control
IWR
D2
+2 h
+6 h
+12 h
+24 h
+48 h
0.0565
I
GATA4 expression
GATA5 expression
ISL1 expression
JUN expression
RARB expression
SOX4 expression
Expression Level
GATA4(+) regulon activity
GATA5(+) regulon activity
ISL1(+) regulon activity
JUN(+) regulon activity
RARB(+) regulon activity
SOX4(+) regulon activity

### Figure S6

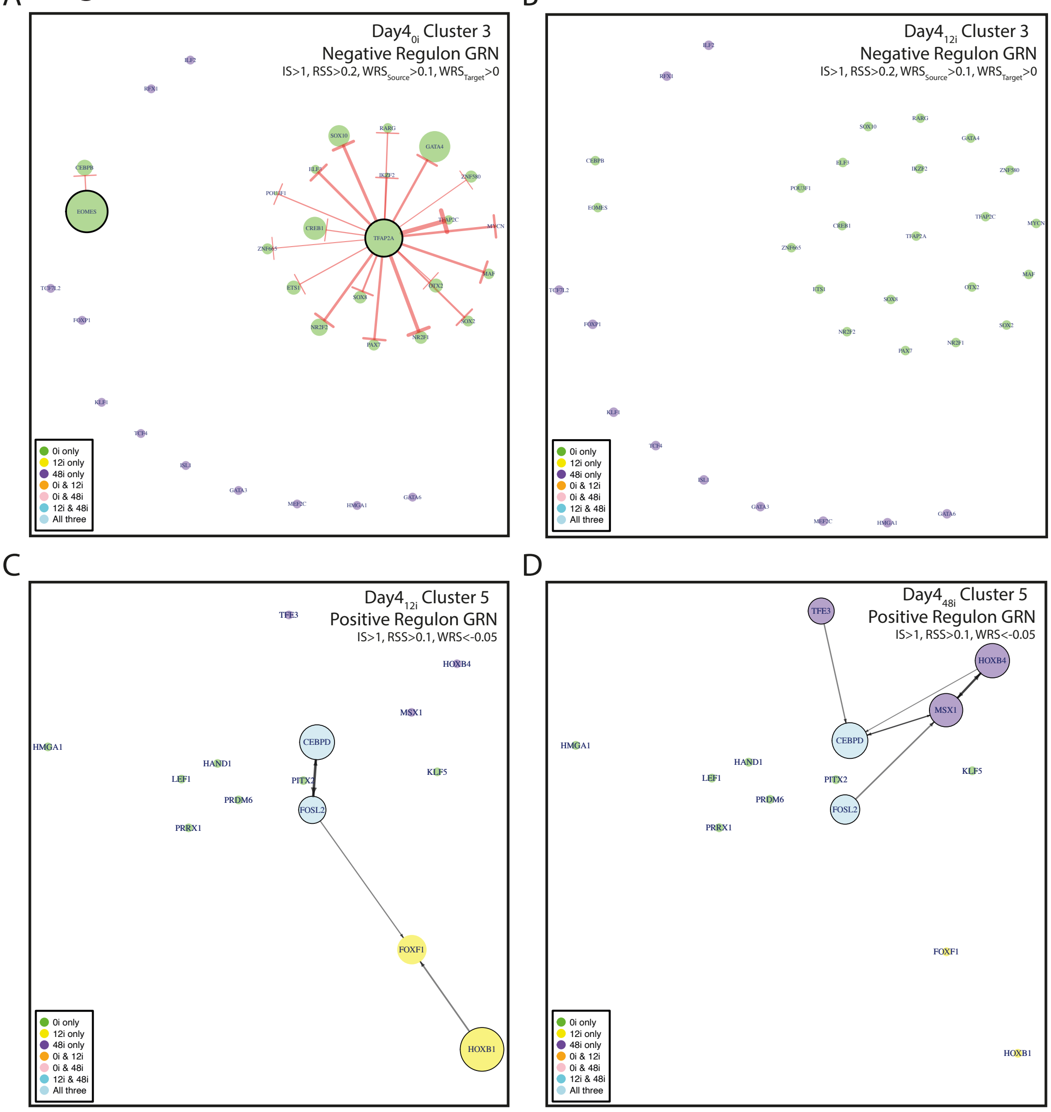
A
Day4 0i Cluster 3
Negative Regulon GRN
IS>1, RSS>0.2, WRS Source >0.1, WRS Target >0
TFAP2A
EOMES
CEBPB
SOX10
RARG
GATA4
ELF3
IKZF2
ZNF580
POU3F1
CREB1
TFAP2C
MYCN
ZNF665
ETS1
SOX8
OTX2
MAF
NR2F2
PAX7
NR2F1
SOX2
TCF7L2
FOXP1
KLF8
TCF4
ISL1
GATA3
MEF2C
HMGA1
GATA6
RFX3
ELF2
B
Day4 12i Cluster 3
Negative Regulon GRN
IS>1, RSS>0.2, WRS Source >0.1, WRS Target >0
C
Day4 12i Cluster 5
Positive Regulon GRN
IS>1, RSS>0.1, WRS<-0.05
TFE3
HOXB4
MSX1
HMGA1
CEBPD
HAND1
LEF1
PITX2
KLF5
PRDM6
FOSL2
PRRX1
FOXF1
HOXB1
D
Day4 48i Cluster 5
Positive Regulon GRN
IS>1, RSS>0.1, WRS<-0.05
0i only
12i only
48i only
0i & 12i
0i & 48i
12i & 48i
All three

**Figure S7**

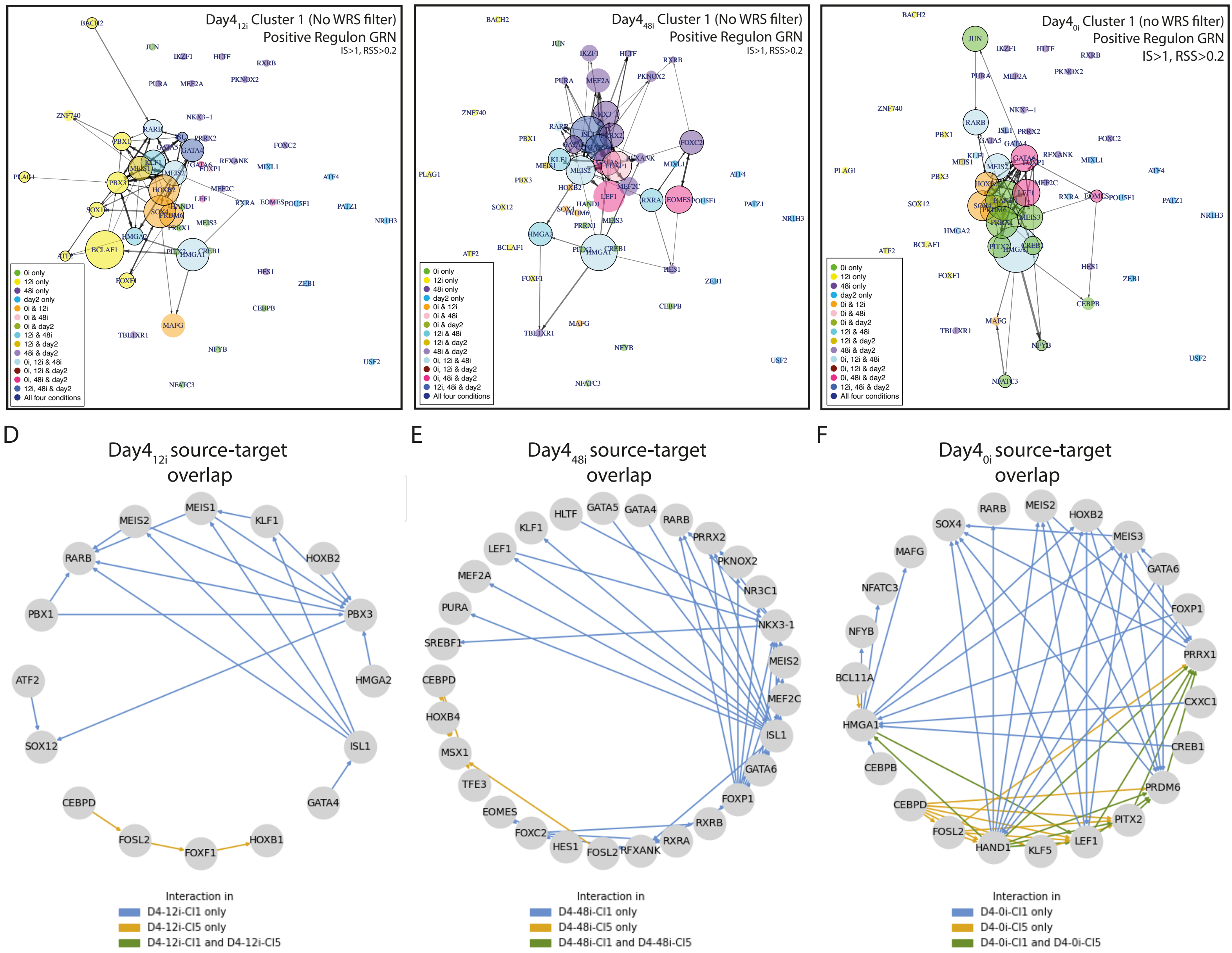
A
Day4 12i Cluster 1 (No WRS filter)
Positive Regulon GRN
IS>1, RSS>0.2
B
Day4 48i Cluster 1 (No WRS filter)
Positive Regulon GRN
IS>1, RSS>0.2
C
Day4 0i Cluster 1 (no WRS filter)
Positive Regulon GRN
IS>1, RSS>0.2
0i only
12i only
48i only
day2 only
0i & 12i
0i & 48i
0i & day2
12i & 48i
12i & day2
48i & day2
0i, 12i & 48i
0i, 12i & day2
0i, 48i & day2
12i, 48i & day2
All four conditions
D
Day4 12i source-target overlap
E
Day4 48i source-target overlap
F
Day4 0i source-target overlap
Interaction in
D4-12i-Cl1 only
D4-12i-Cl5 only
D4-12i-Cl1 and D4-12i-Cl5
D4-48i-Cl1 only
D4-48i-Cl5 only
D4-48i-Cl1 and D4-48i-Cl5
D4-0i-Cl1 only
D4-0i-Cl5 only
D4-0i-Cl1 and D4-0i-Cl5
